# OxPhos in adipose tissue macrophages regulated by BTK enhances their M2-like phenotype and confers a systemic immunometabolic benefit in obesity

**DOI:** 10.1101/2023.10.09.561199

**Authors:** Gareth S. D. Purvis, Massimo Collino, Andrea D. van Dam, Giacomo Einaudi, Yujung Ng, Mayooran Shanmuganathan, Oxford Acute Myocardial Infarction (OxAMI) Study, Smita Y. Patel, Christoph Thiemermann, Keith M. Channon, David R. Greaves

## Abstract

Bruton’s tyrosine kinase (BTK) is a non-receptor bound kinase involved in pro-inflammatory signalling in activated macrophages, however, its role within adipose tissue macrophages remains unclear. We have demonstrated that BTK signalling regulates macrophage M2-like polarisation state by up-regulating subunits of mitochondrially encoded electron transport chain Complex I (*ND4* and *NDL4*) and Complex IV (*mt-CO1*, *mt-CO2* and *mt-CO3*) resulting in an enhanced rate of oxidative phosphorylation (OxPhos) in an NF-κB independent manner. Critically, BTK expression is elevated in adipose tissue macrophages from obese individuals with diabetes, while key mitochondrial genes (mtC01, mtC02 and mtC03) are decreased in inflammatory myeloid cells from obese individuals. Inhibition of BTK signalling either globally (Xid mice) or in myeloid cells (LysMCreBTK), or therapeutically (Acalabrutinib) protects HFD-fed mice from developing glycaemic dysregulation by improving signalling through the IRS1/Akt/GSK3b pathway. The beneficial effects of acalabrutinib treatment are lost in macrophage ablated mice. Inhibition of BTK signalling in myeloid cells but not B-cells, induced a phenotypic switch in adipose tissue macrophages from a pro-inflammatory M1-state to a pro-resolution M2-like phenotype, by shifting macrophage metabolism towards OxPhos. This reduces both local and systemic inflammation and protects mice from the immunometabolic consequences of obesity. Therefore, in BTK we have identified a macrophage specific, druggable target that can regulate adipose tissue polarisation and cellular metabolism that can confer systematic benefit in metabolic syndrome.

**Article high lights:** Obesity and diabetes are associated with inflammation, particularly within the adipose tissue. We have found a new druggable target called Bruton’s tyrosine kinase (BTK) that is highly expressed in adipose tissue macrophages. When BTK is inhibited in macrophages, it allows these cells to undergo a phenotypic switch towards an M2-like pro-resolution macrophage. This achieved by increasing expression of key mitochondrially encoded components of the electron transport chain allowing for enhanced OxPhos. Inhibition of BTK signalling in myeloid cells but not B-cells protects HFD-fed mice from developing glycaemic dysregulation.

## Introduction

White adipose tissue (WAT) can rapidly adapt to nutrient excess through adipocyte hypertrophy and hyperplasia. Expansion of WAT in obesity is associated with the expansion of the stromal vascular compartment, which includes an increase in the number of immune cells of which myeloid cells, including macrophages, are the most abundant ^1^. Obesity is associated with a phenotypic switch of macrophages from homeostatic tissue resident (M2-like) to the recruitment of pro-inflammatory (M1-like) macrophages ^2^. M2-like adipose tissue macrophages (ATM) have the capacity for metabolic adaption under nutrient surplus, unlike their pro-inflammatory M1-like counterparts ^3^; in obesity ATM have reduced ability to adapt to changes in excessive nutrient demand ^4^. Increased inflammation and increased numbers of ATM, in particular, result in the development of systemic insulin resistance^5^.

Macrophage energy metabolism is intrinsically linked to their polarisation state. Pro-inflammatory macrophages have enhanced glycolytic metabolism, pentose phosphate pathway and fatty acid synthesis but have breaks in the TCA cycle leading to reduced mitochondrial oxidative phosphorylation (OxPhos) ^9^. M2/M(IL-4) macrophages also utilise glycolytic metabolism, increase fatty-acid oxidation and have an enhanced ability to utilise OxPhos, even in the absence of glycolysis by utilising glutamine ^10^. *In vitro* experiments suggest that phenotypic switching from M(LPS/IFN γ) to M(IL-4) macrophages can only be achieved by over-coming the mitochondrial dysfunction caused by excessive NO production, re-enforcing the central role of OxPhos in M(IL-4) polarisation^11^.

Bruton’s tyrosine kinase is best known for its critical role in B-cell development. Indeed, patients with point mutations within the *Btk* gene have X-linked agammaglobulinemia (XLA) which is a genetic immunodeficiency characterised by the absence of mature B-cells. However, BTK is also highly expressed in myeloid cells, where its functions are less well understood ^12,13,14^. BTK plays a central role in macrophage pro-inflammatory signalling regulating the production of cytokines and chemokines ^15,16^, and regulating myeloid cell recruitment ^17,18^.

We, and others, have reported that both genetic and pharmacological inhibition of BTK reduces the activation of NF-kB and the NLRP3 inflammasome in pre-clinical models of obesity/T2D ^19^, polymicrobial sepsis ^20,21^ and cerebral ischaemia ^22^. However, the role of BTK in the polarisation of macrophages in obesity-derived inflammation is unknown, and the role of macrophages in regulating WAT inflammation remains unclear. In this study we will investigate at a molecular level how BTK signalling alters macrophage polarisation and metabolic state within adipose tissue, and how this affects glycaemic regulation of mice following HFD feeding.

## Methods

### Human Studies Ethics

Ethics approval was obtained (HRA NRES Research Ethics Reference Number 11/SC0397 and 12/SC/0044). Patients with sequence confirmed XLA provided written consent prior to blood sampling. Age and sex matched healthy donors (HD) provided written consent prior to blood sampling; any individuals with any significant past medical history or on regular medications were excluded.

### Human Monocyte-Derived Monocyte (hMDMo) and Macrophages (hMDMs)

Peripheral blood mononuclear cells (PBMCs) were purified from whole blood collected into EDTA tubes by density centrifugation over Histopaque (Sigma). The PBMC layer was carefully harvested and then washed repeatedly with PBS and centrifugation. CD14 ^+^ monocytes were then isolated from the PBMCs by positive selection using magnetic beads conjugated to anti-CD14 antibody (Miltenyi Biotec, Bergisch Gladbach, Germany).

Monocytes were then ready for experimentation. To generate human monocyte derived macrophages monocytes were maintained in RPMI 1640 medium supplemented with heat inactivated 10% FBS, 50 ng/ml macrophage colony-stimulating factor (M-CSF, BioLegend, San Diego, CA, USA), 50 U/ml penicillin/streptomycin for 7 days.

### Animal Studies Ethics

All animal experiments were conducting in accordance to the Animal (Scientific Procedures) Act 1986, with procedures reviewed by the Local ethical review body (AWERB) and conducted under project license P144E44F2 at the University of Oxford. CBA/N (XID) mice containing a single amino acid change (R28C) conferring X-linked immunodeficiency (*Xid)* to the strain ^23^ were originally purchased from JAX Lab. CBA/CaCrl (WT) were purchased from Charles River and served as wild type (WT) control for XID mice. BTK floxed and CD19 Cre mice were kindly provided by Dr Wasif Khan and LysM Cre mice were originally purchased from JAX Lab.

## Cell Culture of Primary macrophages

### Murine Bone Marrow-Derived Macrophages (BMDMs)

BMDM were generated as previously described ^18^. Briefly, fresh bone marrow cells from tibiae and femurs of male CBA, XID mice were isolated and cultured in Dulbecco’s Modified Eagle’s medium (DMEM) supplemented with 10% heat-inactivated fetal bovine serum (FBS), 10% L929 cell-conditioned media as a source of macrophage colony-stimulating factor, and 50 U/ml penicillin/streptomycin for 7 days. Bone marrow cells were seeded into 8 ml of medium in 100 mm non-tissue culture treated Petri dishes (Thermo Fisher Scientific, Sterilin, UK). On day 5, an additional 5 ml of medium was added. Gentle scrapping was used to lift cells. BMDMs were then counted and resuspended in DMEM containing 2 % FBS and 1 % penicillin/streptomycin media at the desired cell concentration for experiments.

### Resident Peritoneal Macrophages

Male CBA, XID mice were sacrificed and peritoneal cavities were lavaged with 10 ml ice-cold PBS supplemented with 5 mM EDTA. Cells were pelleted by centrifugation, resuspended in DMEM 10% FBS, and plated for 2–4 h to allow macrophages to attach to the plate.

### Cell Culture of Cell lines

THP-1 cells were grown to a maximal confluence of 2×10 ^6^ per mL in T-175 culture flasks in RMPI containing heat-inactivated 10% FBS, 1 % penicillin/streptomycin 37°C in 5% CO2. Cells were passaged every 4 days, and used until a maximum passage of 12. THP-1 cells were then plated for experiments or differentiated into macrophages by adding phorbol 12-myristate-13-acetate (PMA) (50 ng/mL) for 48 h followed by 24 h recovery without PMA.

### RNA Seq analysis

Bone marrow derived macrophages were generated as previously described ^24^. Following BMDM stimulation, total RNA was extracted from each sample using the RNeasy kits (QIAGEN) with on column DNA digestion. RNA was quantified using a ND-1000 spectrophotometer. RNA quality control was performed using an Aglient Tape Station to confirm RNA integrity and concentration. Libraries were sequenced together on a single Illumina HiSeq 4000 lane at the Oxford Genomics Centre using a 28×98bp paired-end read configuration. Base-calling and demultiplexing were performed using “cellranger mkfastq” pipeline in order to produce Cell Ranger compatible FastQ files. FastQ files were quality controlled using FastQC (Andrews et al., 2010). Data were aligned against the pre-built mouse reference GRCm38 (Ensembl 93) primary assembly and annotation files. Cell Ranger’s “count” pipeline performs the alignment, filtering and barcode counting. Pathway enrichment analysis was undertake using Ingenunity Pathways Analysis (IPA) software (Qiagen) using the top 500 differentially expressed genes.

### Seahorse XFe96 analysis of mitochondrial function

On day 7 of culture macrophages were plated at a density of 7.5 × 10 ^4^ cells per well into XFe96 microplates. Cells were left to attach for 2 hours before stimulation with LPS (100 ng/mL) and IFNγ (20 ng/mL) or IL-4 (20 ng/mL) if required, for 18 hours. Extracellular flux analysis was then performed. One hour prior to the assay, cells were washed and the culture medium was replaced with Seahorse XF Base Medium (modified DMEM without Phenol Red, pH 7.4; Agilent), supplemented with glucose (10 mM, Sigma), glutamine (2 mM, Sigma), and sodium pyruvate (2 mM, Sigma), before being incubated at 37°C, at atmospheric CO2 levels. Oxygen consumption rate (OCR) and extracellular acidification rate (ECAR) were measured using the XFe96 analyser (Agilent). 6 base-line OCR measurements were taken followed by 3 measurements after sequential injection of the following compounds 2 μM, oligomycin (Sigma), 2 μM FCCP (Sigma), and combined 0.5 μM antimycin A (Sigma), and 0.5 μM rotenone (Sigma). Cells were measured in 4 replicate wells and all Seahorse was data normalized to cell number. For calculations the third reading in each case was used as the most stable point, except for FCCP were the highest reading was taken.

### Quantitative real-time RT-PCR

RNA was extracted with TRIzol reagent (Thermo Fisher Scientific), and purified using Direct-zol RNA kit; total RNA concentration and quality was determined with a ND-1000 spectrophotometer (Nano Drop Technologies, Wilmington, DE, USA). cDNA was synthesized from 1,000 ng total RNA using the QuantiTect Reverse Transcription kit (Qiagen, Manchester, UK) according to the manufacturer’s instructions. Real-time quantitative PCR was performed using Sybr Select gene expression master mix (Life Technologies) in the StepOnePlusTM thermal cycler (Applied Biosystems). Primers were purchased from Qiagen *(mt-CO1, mt-CO2, mt-CO3, 18S, cxcl1, ccl2, tnfa, Il-6, Il-1b*). Cycle threshold values were determined by the StepOne software and target gene expression was normalized to housekeeping gene (*18S*). Relative expression results were plotted as mRNA expression divided by housekeeping gene.

### Western blotting

Cells were lysed by adding RIPA buffer (Sigma-Aldrich) supplemented with protease and phosphatase inhibitors (Sigma-Aldrich) followed by manual disruption. Protein concentration was determined by using a BCA protein assay kit (Thermo Fisher Scientific). Total cell protein (20–30 mg) was added NuPAGE LDS sample Buffer (4X) containing NuPAGE Sample Reducing Agent (10X). Samples were then resolved on Bis-Tris gels and electrotransferred onto polyvinyldenedifluoride (PVDF) membranes using the Mini Gel Tank and the Mini Blot Module (ThermoFisher Scientific). Membranes were blocked with 5% milk in Tris-buffered saline with Tween (TBS-T) (Tris-buffered saline, 0.1% Tween-20) for 1 h at RT and then incubated with the primary antibody (mt-CO1, mt-CO2, or Total OXPHOS Rodent WB Antibody Cocktail) in 5% BSA/TBS-T overnight at 4°C. Next, membranes were incubated with an HRP-conjugated antibody for 1 h at RT. Protein bands were visualized by incubating the membranes for 5 min with Amersham ECL prime and subsequent exposure to X-ray film over a range of exposure times. GAPDH or B-actin were used as loading controls. For successive antibody incubations using the same membrane bound antibodies were removed with stripping buffer (Thermo Fisher Scientific). Liver and skeletal muscle extracts were prepared as previously ^26^. Briefly, tissues weighting 50 mg were homogenized and centrifuged at 10,000 g for 20 min at 4 C. Supernatants were collected and the protein content was determined using a BCA protein assay (Pierce Biotechnology Inc., Rockford, IL, USA). Equal amounts of total protein extracts were added with BIORAD Laemlli Sample Buffer 4x with 0.5% DTT, separated by sodium dodecyl sulphate-polyacrylamide gel, electrotransferred to a PVDF membranes and blocked using 10% milk solution in TBS-T for 1h at room temperature. Membranes were probed overnight at 4° C with rabbit anti-Ser ^307^ IRS-1 (#2381), mouse anti-total IRS-1 (#3194), rabbit anti-Ser ^473^ AKT (#4060), rabbit anti-total AKT (#9272), rabbit anti-Ser ^9^ GSK-3β (#9332) and rabbit anti-total GSK-3β (9315) and rabbit anti-B actin (#4970) used as loading control. The blots were then incubated with secondary antibodies conjugated with horseradish peroxidase (dilution 1:10000), detected with Clarity Western ECL substrate (BioRad, California, USA) and quantified by densitometry using analytic software (Quantity-One, Bio-Rad) and the results were normalised to the controls.

### scRNA-seq data resources and analysis

The present study used the 10x scRNA-seq dataset published by Hildreth *et al* ^27^. Briefly, human subcutaneous WAT samples were obtained from 6 healthy females, *i.e.* without history of cardiovascular disease, liver disease, diabetes or immunological disorders. Three donors were lean (BMI 20.5, 24.8, 16.9 and three were obese (BMI 33.4, 30, 31.6). Donors were between 32-43 years old. Adipose tissue biopsies were homogenised, collagenase digested and centrifuged to isolate the stromal vascular fraction (SVF). SVF samples were sorted into nonimmune cells (CD45^-^) and immune cells (CD45^+^) before generation of scRNA-seq libraries (using 10x Genomics Chromium) and sequencing. Upon alignment and generation of the expression matrix, the R package Seurat was used for quality control, data normalization and scaling, selection of cells for further analysis and clustering and annotation of cell types, of which the details are described in the original paper. This information was stored in an expression matrix and a metadata table, which we obtained from the O’Sullivan lab. The data were further analysed and visualised in R v4.0.3 using Seurat v4.0.5 (REF https://satijalab.org/seurat) visualization tools for UMAP and violin plots. The metadata was used to subset the complete SVF samples (61311 cells) from the original dataset, excluding any other FACS-sorted immune cell populations that were included in the original paper. To calculate sample composition, in each sample the number of cells of one specific type was divided by the total number of cells in the corresponding sample. The resulting fractions were expressed as a percentage. Murine single cells transcriptomics analysis was conducted using the online open resource (https://hastylab.shinyapps.io/MAIseq/).

### Diet-induced obesity model

In this model, male mice (CBA/CaCrl - Wild type or CBA/N - XID) were randomised to receive a defined ‘chow’ diet [LabDiet®, St. Louis (5053: protein provides 25%, fat 13%, and carbohydrate 62% of the total calories)] or high fat diet [TestDiet®, St. Louis (D12331: Red diet; protein provides 17%, fat 58%, and carbohydrate 25% of the total calories)] with free access to water for 12 weeks.

#### Cell type specific knock out study

Heterozygous male CD19 Cre (CD19 ^+/-^) and LysM Cre (LysM^+/-^) mice were crossed with female BTK floxed mice, all male mice born are hemizygous for the BTK floxed allele. The BTK floxed allele was confirmed using the following primers Ex7F1: 5’ CCTACCCCAGAGGAAGATCAG and I7/8R2: 5’ CTCACTCTGTAGACCAGGTAGGCCT. CD19Cre mice were genotyped using the following primers CD19.8: 5’ AAT GTT GTG CTG CCa TGC TGC CTC CD19.9: 5’ GTC TGA AGC ATT CCA CCG GAA and CD19CreRev 5’ TTC TTG CGA ACCT CA TCA C as previously described ^28^. LysMCre mice were genotyped using genotyping protocol provided by supplier. Mice were fed the same high fat diet with free access to water for 12 weeks.

#### Therapeutic study

Male WT C57BL/6 mice were fed the same high fat diet with free access to water for 12 weeks, after the initial 6 weeks mice were randomly assigned to receive either vehicle (5 % DMSO in 30 % w/v 2-hydroxypropyl-β-cyclodextrin) or acalabrutinib (1 mg/kg; p.o.; by oral gavage) 5 times per week for a further 6 weeks. Further details are included in the Supplementary material.

#### Macrophage ablation study

Male WT C57BL/6 mice were fed the same high fat diet with free access to water for 10 weeks, after the initial 5 weeks mice were randomly assigned to a receive either control liposome (i.p.; weekly; Liposoma Research, USA) or clodronate loaded liposome (0.5 mg/kg; i.p.; weekly; Liposoma Research, USA) co-administered with vehicle or acalabrutinib (1 mg/kg; p.o.; by oral gavage; 5 times per week) for a further 5 weeks. Body weight, food intake, and calorie intake were measured weekly throughout all the experiment to monitor feeding behaviour, no observable differences were seen between any group.

#### Oral Glucose Tolerance Test (OGTT)

Mice were fasted for 6 h with free access to water. At the end of the 6 h fasting, the body weight and fasting blood glucose were measured using tail vein blood. Mice then received an oral bolus of glucose (2 g/kg; dissolved in drinking water) via oral gavage. Blood glucose levels were then measured at 15, 30, 45, 60, 90, and 120 min post glucose administration using blood glucose meter Accu-Chek® (Accu-Chek Compact System; Roche Diagnostics, Basel, Switzerland).

#### Insulin Tolerance Test (ITT)

Mice were fasted for 4 h with free access to water. At the end of the 4 h fasting, the body weight and fasting blood glucose were measured using tail vein blood. Mice then received a dose of insulin (0.75 U/kg; i.p.). Blood glucose level was then measured at 15, 30, 45, 60, 90, and 120 min post insulin administration using blood glucose meter Accu-Chek® (Accu-Chek Compact System; Roche Diagnostics, Basel, Switzerland).

#### Biochemical analysis blood

Blood was taken following terminal anaesthesia. Biochemical analyses of plasma lipids were performed on heparinized blood plasma using enzymatic assays by a commercial laboratory (MRC Harwell, Oxford, UK).

### Tissue processing and histology

Organs were fixed with 4% paraformaldehyde solution (Sigma-Aldrich) immediately and subjected to paraffin embedding after a minimum of 1 week’s fixation. After deparaffinization, 5 μm‒thick sections were stained with hematoxylin and eosin (Sigma-Aldrich) for general morphology and adipocyte size measurement. Adipocyte size was measured using the ImageJ.

### Stroma vascular fraction isolation

Fat was washed in 0.1% bovine serum albumin in PBS, finely minced, and enzymatically digested with 2 mg/mL type I collagenase (type I collagenase) in HBSS for 40 min at 37 °C. After digestion, the tissue-collagenase mixture was centrifuged at 100g for 8 mins, buoyant adipocytes were removed, the remaining supernatant and cell pellet was centrifuged at 400 g for 5 mins to collect SVF.

### Flow cytometry

Cells were washed in FACS buffer (0.05 % BSA, 2 mM EDTA in PBS pH 7.4), blocked using anti-CD16/32 for 15 mins at 4°C, followed by antibody staining for surface markers with appropriate isotype controls for 30 mins at 4°C. Fluorescence minus one (FMO) controls were used for gating. For intracellular pBTK staining, cells were cell surface marker stained followed by fixation and permeabilisation then stained for intercellular pBTK for 60 mins at room temperature. Data were acquired using BD Fortessa X20 cytometer and Diva software (BD Biosciences) and then analysed using FlowJo (Tree Star Inc, USA) software.

### Statistical analysis

Animal experiments were designed, where possible, to generate groups of equal size. Power calculations were used to calculate the group size based on an expected effect size of 30%. Where possible, blinding and randomisation protocols were used. All data in the text and figures are presented as mean ± standard error mean (SEM) of *n* observations, where *n* represents the number of animals studied (*in vivo*) or independent values, not technical replicates (*in vitro*). For western blot data, all data are represented as fold change to mean control (normal chow diet). Statistical analysis was only undertaken for studies where each group size was at least *n* = 4. For western blot analysis, some representative data are shown where group size is less then *n* = 4. When the mean of two experimental groups were compared, a two-tailed Students *t*-test was performed. Normally distributed data without repeated measurements were assessed by a one-way ANOVA followed by Bonferroni correction if the *F* value reached significance. In all cases a *P* < 0.05 was deemed significant. All statistical analysis was calculated using GraphPad Prism 7 for Mac (GraphPad Software, San Diego, California, USA; RRID:SCR_002798).

### Data and resource availability statements

The datasets generated during and/or analysed during the current study are available from the corresponding author upon reasonable request. No applicable resources were generated or analysed during the current study.

## Results

### Bruton’s tyrosine kinase regulates macrophage polarisation

We first assessed BTK activation during polarisation of human macrophages. THP-1 cells were treated with PMA to induce differentiation to into macrophages and subsequently stimulated with LPS/INFy or IL-4. Compared to M0, M(LPS/IFN γ) human macrophages had increased Tyr ^223^ phosphorylation on BTK, while there was no difference following IL-4 stimulation (Figure 1A; Supplementary Figure 1A). Specifically, HLA-DR ^hi^ THP-1 cells that were stimulated with LPS/INF γ had significant increase in Tyr ^223^ phosphorylation on BTK (Figure 1B), while CD206 ^hi^ THP-1 cells that were stimulated with IL-4 had a significant decrease in Tyr ^223^ phosphorylation on BTK (Figure 1C). These results demonstrate that BTK activation (phosphorylation) is linked to human macrophage polarisation state.

**Figure 1:**
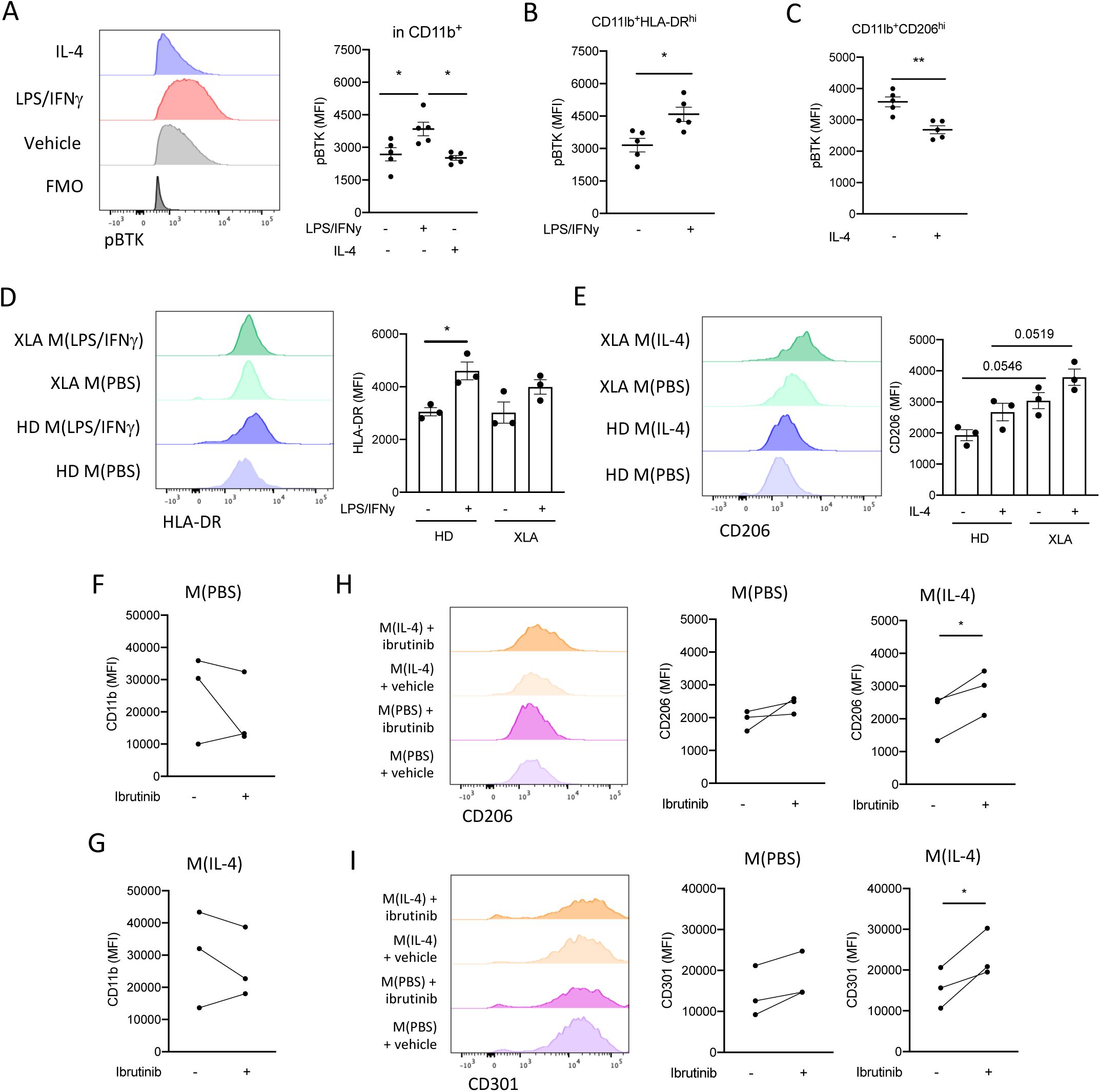
Bruton’s tyrosine kinase regulates human and murine macrophages polarisation. (A) Representative histograph of mean fluorescent intensity (MFI) of pBTK on THP-1 cells and quantification. (B) pBTK (MFI) in CD11b ^+^ HLA-D R^hi^ THP1 cells treated with LPS/IFN γ or vehicle. (C) pBTK (MFI) CD11b ^+^CD206^hi^ THP1 cells treated with IL-4 or vehicle. (D) representative histograph of the mean fluorescent intensity (MFI) of HLA-DR on human monocyte derived macrophages generated form healthy donors and patients with XLA stimulated with LPS/IFN γ or PBS. E) Representative histograph of the mean fluorescent intensity (MFI) of CD206 on human monocyte derived macrophages generated form healthy donors and patients with XLA stimulated with IL-4 or PBS. (F-G) Quantification of mean fluorescent intensity (MFI) of CD11b on human monocyte derived macrophages form healthy donors treated stimulated with IL-4 or PBS in the presence of ibrutinib or vehicle. (H) Representative histograph and quantification of mean fluorescent intensity (MFI) of CD206 on human monocyte derived macrophages form healthy donors treated stimulated with IL-4 or PBS in the presence of ibrutinib or vehicle. (I) Representative histograph and quantification of mean fluorescent intensity (MFI) of CD301 on human monocyte derived macrophages form healthy donors treated stimulated with IL-4 or PBS in the presence of ibrutinib or vehicle. Data shown are means ± SEM of n biological replicates. *P< 0.05 **P< 0.01; a student’s t-test was performed when there were two variables, and a one-way ANOVA was performed with Bonferroni post hoc test when there were more than two variables.

Patients with X-linked immunodeficiency (XLA) have point mutations within the *Btk* gene, causing loss of signalling function ^29^. We collected whole blood samples from patients with XLA or healthy donors (HD) and isolated monocytes to generate monocyte-derived macrophages (MoDM). As expected, addition of LPS/IFN γ to hMoDM resulted in an increased expression of HLA-DR, and there was no difference between in HLA-DR expression between HD and XLA patients (Figure 1D). However, when compared to M(IL-4) from HD, M(IL-4) from XLA patients had increased expression of CD206 (Figure 1E). These results are consistent with BTK signalling supressing the M2-like phenotype in human macrophages.

We next investigated whether pharmacological inhibition of BTK, using the FDA approved BTK inhibitor Ibrutinib, would result in the same phenotype observed in hMoDM from XLA patients. When compared to vehicle-treated macrophages, Ibrutinib-treated macrophages displayed no difference in CD11b expression regardless of their polarisation state (Figure 1F/G), but M(IL-4) cells treated with ibrutinib had significantly increased expression of CD206 (Figure 1G) and CD301 (Figure 1I), recapitulating the results seen in macrophages from XLA patients. Collectively, these results indicate that BTK signalling modulates macrophage polarisation state in human macrophages. Specifically, inhibiting BTK signalling allows macrophages to undergo a phenotypic switch towards an M2-like state.

### Differential gene expression data from WT and XID M(IL-4) BMDM reveals a central role for Oxidative Phosphorylation (OxPhos)

Murine macrophages from XID mice, which have a point mutation in the *Btk* gene, displayed a similar phenotype to human patients with XLA; enhanced cell surface expression of CD206, which was further enhanced following IL-4 stimulation (Figure 2A/C). Accordingly, we next used RNA Sequencing to assess gene expression profiles of WT and XID M(IL-4) BMDMs to understand the mechanisms behind the enhanced M2-like phenotype in macrophages from XID mice. In XID M(IL-4) BMDMs 76 genes showed differential expression (DE) with a >5 % False Discovery Rate (Figure 2D). These included *lL-4RA, Relna, Arg1* and *Irf4* (Figure 2E). Ingenuity Pathway Analysis (IPA) of the DE genes revealed significant enrichment in pathways key to macrophage functions. These included pathways involved in protein translation (EIF2 signalling), energy metabolism (mTOR signalling and oxidative phosphorylation), phagocytosis and cellular motility (PI3K/Akt signalling and Rac signalling) (Supplementary Figure 2A).

**Figure 2:**
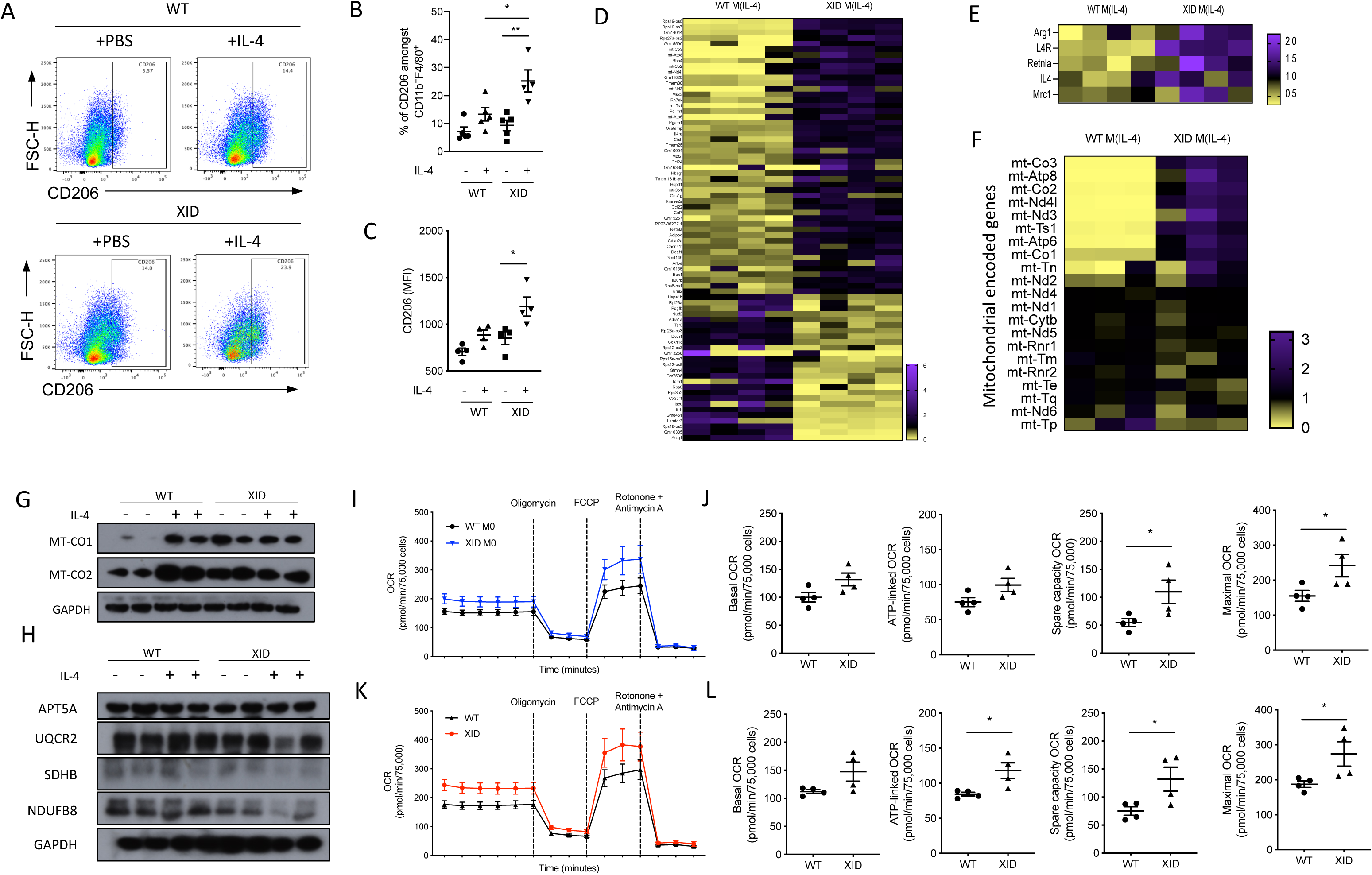
M(IL-4) macrophages from XID mice have enhanced ability to utilise oxidative phosphorylation. (A) Bone marrow derived macrophages generated from WT and XID mice were polarised with IL-4. Representative flow cytometry plots of BMDM (CD11b ^+^ F4/80^+^) showing expression of CD206 (B) and quantified MFI (C). RNA sequencing was performed on M(IL-4) BMDM generated from WT and XID mice. (D) Heatmap showing differentially expressed genes. (E) Heatmap of macrophage M2 related genes. (F)Heat map of mitochondrially encode genes. (G) Representative western blots of mitochondrial proteins mt-CO1 and mt-CO2 in M0 and M (IL-4) BMDM generated from WT and XID mice.(H) Representative western blots of mitochondrial proteins from Complex I (NDUFB8), Complex II (SDHB), Complex III (UQCR2) and Complex V (APT5A) in M0 and M(IL-4) BMDM generated from WT and XID mice. (F-I) Oxygen consumption rate (OCR) was measured using XFe 96 Seahorse bioanalyzer with compounds used to determine basal, ATP-linked, spare capacity and maximum in MO BMDM (I/J) (black solid line -WT; blue solid line - XID) and in (K/L) M(IL-4) BMDM (black solid line - WT; red solid line - XID). Data shown are means ± SEM of n biological replicates. *P< 0.05; a student’s t-test was performed when there were two variables, and a one-way ANOVA was performed with Bonferroni post hoc test when there were more than two variables.

### BTK signalling regulated mitochondrially encoded genes in M(IL-4) BMDM

Energy metabolism from the electron transport chain is essential for M(IL-4) polarisation ^30^. IPA of DE genes revealed oxidative phosphorylation (OxPhos) to be significantly up-regulated pathway in M(IL-4) BMDMs from XID mice. Importantly, only genes from the light strand of the mitochondrial genome were significantly up-regulated including transcripts for complex I subunits (*ND3* and *ND4L*), complex IV subunits (*mt-C01*, *mt-C02* and *mt-C03*) and complex V subunits (*Atp6* and *Atp8*) (Figure 2F). Western blot analysis confirmed increased protein levels of mtCO1 and mtCO2 in M(IL-4) BMDM compared to WT M0 BMDM (Figure 2G). Furthermore, there was no difference in the levels of other mitochondrial proteins which are not encoded on the mitochondrial genome NDUFB8 (from complex I), SDHB (from complex II), UQCR2 (from complex III) and ATP5A (from complex V) (Figure 2H).

We next investigated if the observed increased expression of electron transport chain proteins altered the rate of OxPhos in BMDM from XID mice, using real time analysis of oxygen consumption rate (OCR). When compared to WT M0 BMDMs, M0 BMDMs from XID mice had enhanced OCR (spare capacity and maximal) (Figure 2I and J), which was further enhanced in M(IL-4) BMDMs from XID mice, as was ATP-linked OCR (Figure 2K and L).

We and others have previously shown that BTK regulates transcription factor NF-κB (p65) activity and nuclear translocation ^31,21,13^, additionally Cogswell *et al*. proposed that NFκ-B is located within the mitochondria and can regulate mt-CO1 and mt-CO2 gene expression ^32^. Therefore, we hypothesised that BTK could be signalling through NF-κB to specifically alter the expression of key mitochondrially encoded genes. Using super resolution immunofluorescence microscopy p65 did not co-localise with the mitochondria in either WT or XID BMDM (Supplementary Figure 3A). Additionally, inhibition of IKK signalling which sequestered p65 in the cytoplasm had no effect in mitochondrially encoded genes *mtC01* and *mtCO3*, or nuclear encoded electron transport component *Cybb (Supplementary* Figure 3B). Therefore, we can conclude that BTK signalling via NF-kB (p65) does not alter mitochondrially encoded gene expression in BMDM.

Another possible mechanism for the enhanced capacity for increased OxPhos is that there is increased number of mitochondria in M(IL-4) BMDM from XID mice. However, no difference in mitochondria content in M(IL-4) macrophages from XID mice was detected (Supplementary Figure2C), but rather the transcription of a subset of mitochondrial encoded genes. Interestingly, these are located on the light strand of the mitochondrial genome suggesting that a mitochondrial specific transcriptional event is occurring rather than increased mitochondrial biogenesis. These data demonstrate that BTK signalling which can alter macrophage ability to utilise OxPhos by regulating mitochondrial genome encoded gene expression, independent of BTK:NF-kB signalling and mitochondrial biogenesis.

### Bruton’s tyrosine kinase is highly expressed in myeloid cells within murine and human adipose tissue

Adipose associated inflammation is a major source of pro-inflammatory signalling that can lead to metabolic syndrome, therefore, we wanted to investigate in which cells BTK is expressed within adipose tissue. Using single cell RNA Sequencing data (GSE182233) immune cells were identified in both lean and obese murine adipose tissue. *Btk* w a s expressed in myeloid cells (macrophages, monocytes, dendritic cells and mast cells) and B-cells; with limited expression is T-cells, ILC’s and stromal cells (Figure 3A). A similar expression pattern of *BTK* was found within human adipose tissue from lean and obese individuals (GSE155960 and GSE156110) with *BTK* expression restricted myeloid cells and B-cells (Figure 3B). No difference was detected in BTK levels between lean and obese individuals. We next wanted to assess if BTK dependent mitochondrial encoded genes were differentially expressed in obesity. As expected mtCO1, mtCO2 and mtCO3 was ubiquitously expressed in all cell populations in the SVF from both murine and human samples (Figure 3C and D). *mtCO1, mtCO2* and *mtCO3* had decreased expression in murine macrophages, lipid associated macrophages and monocytes from HFD fed mice (Figure 3E). Similarly, within human adipose there was a significant decrease in the expression of *mt-CO1*, *mt-CO2* and *mt-CO3* in macrophages, and classical monocytes from obese individuals (Figure 3F). However, *mt-CO1, mt-CO2* and *mt-CO3* expression was higher in non-classical monocytes in obese individuals (Figure 3F). Collectively these data suggest that within pro-inflammatory myeloid cells expression of key components of Complex IV are down regulated.

**Figure 3:**
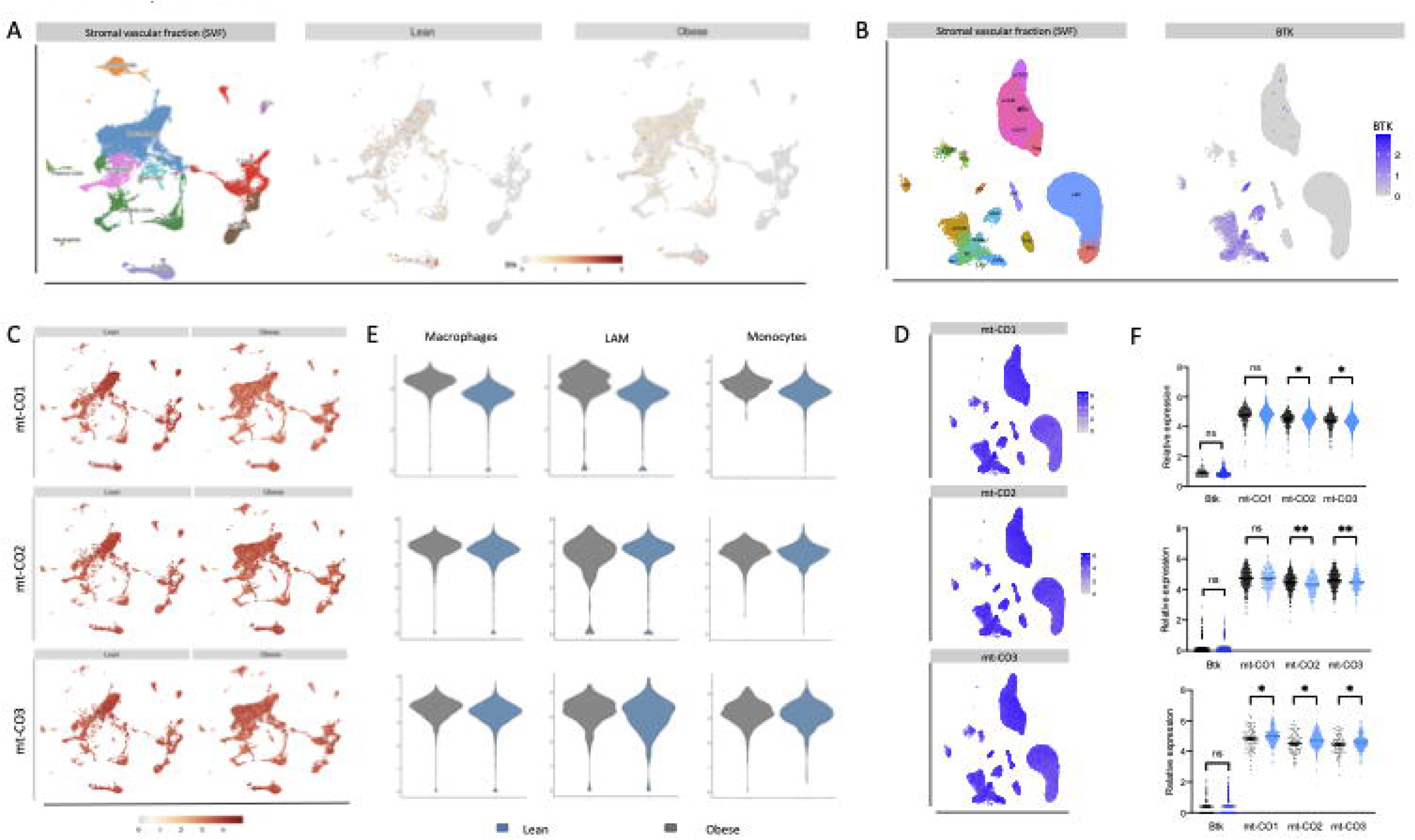
BTK is highly expressed in myeloid cells isolated from human adipose tissue. (A) uMAP of the stromal vascular fraction (SVF) isolated from adipose tissue from lean and obese mice initially coloured by cluster identified from Louvain clustering. Relative *BTK* expression is represented in lean and obese mice separately. Relative *Btk* expression is represented in lean and obese mice separately B) uMAP plots of the stromal vascular fraction (SVF) isolated from adipose tissue from lean and obese human individuals coloured by cluster identified from Louvain clustering. Relative *Btk* expression is represented in combined cells from lean and obese mice. (C) uMAP visualisation of *mt-CO1*, *mt-CO2* and *mt-C03* in both lean and obese mice. (D) Combined uMAP visualisation of *mt-CO1*, *mt-CO2* and *mt-C03* in human individuals. (E) Relative expression of *mt-CO1*, *mt-CO2* and *mt-C03* in macrophages, lipid associated macrophages (LAM) and monocytes. (F) Relative expression of *mt-CO1*, *mt-CO2* and *mt-C03* in inflammatory macrophage (top), classical monocytes (middle) and non-classical monocytes (bottom).

### HFD-fed XID mice are protected from development of diet induced metabolic syndrome

Activation of ATM is associated with increased risk of developing glycaemic dysfunction, while feeding mice a HFD leads to increased ATM number and activation ^33^. We therefore wanted to critically assess the role of BTK signalling in diet induced obesity and metabolic syndrome. WT and XID mice fed a HFD do not differ in weight (Figure 4A-C). Lipid analysis of blood plasma samples revealed that XID mice fed a HFD had no overall difference in total cholesterol levels (Figure 4D), however had significantly lower TG levels (Figure 4E) and significantly increased HDL levels (Figure 4F) compared to WT mice fed a HFD; demonstrating an improvement in blood lipid profiles.

**Figure 4:**
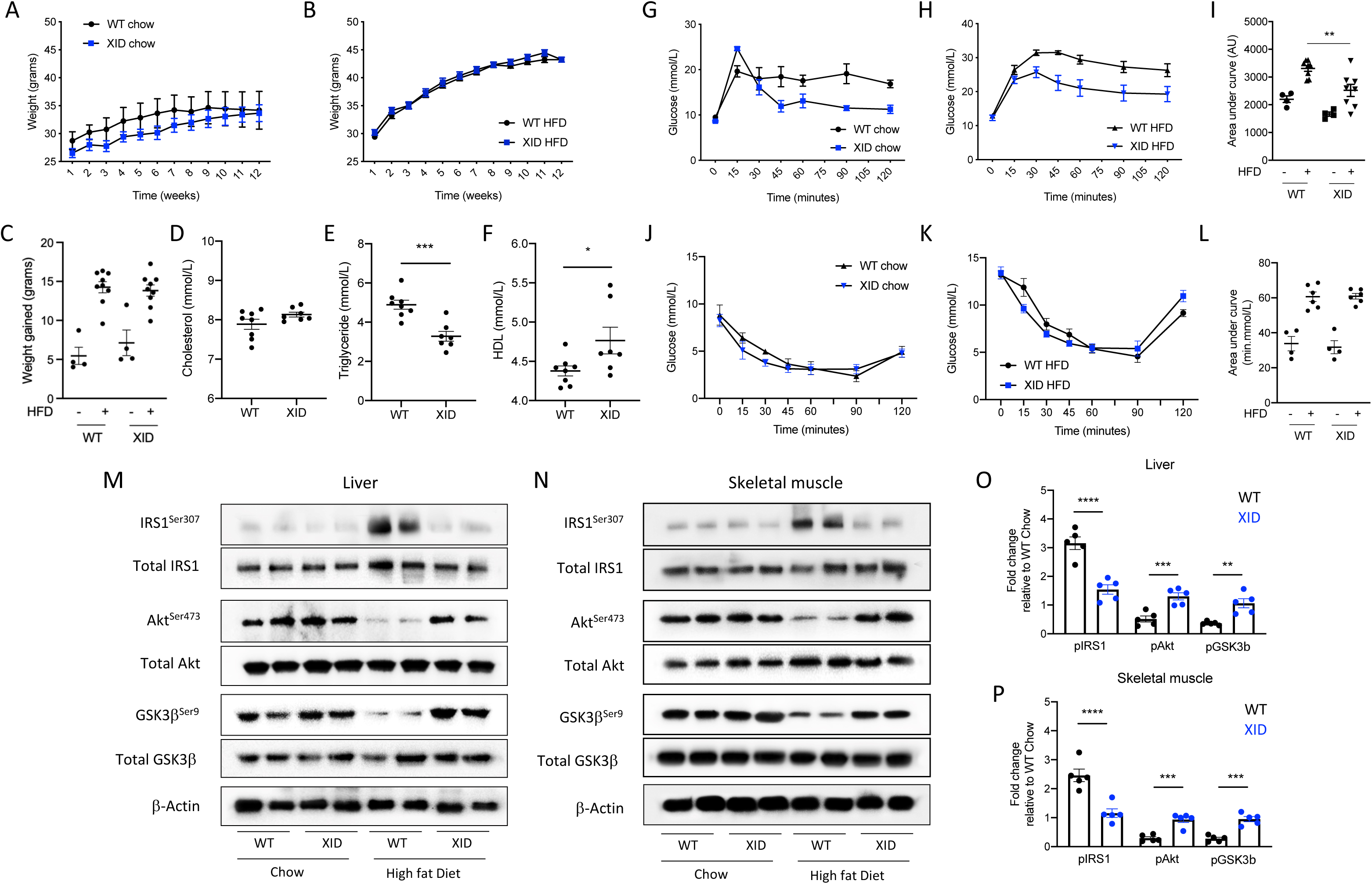
XID mice are protected against the development of HFD diet induced glycaemic dysregulation. WT and XID mice were fed a chow or HFD 12 weeks and body weight was measure each week (A) chow-fed mice and (B) HFD-fed mice and summarised as weight gained (C). Terminal blood plasma analysis of (D) cholesterol, (E) triglyceride and (E) HDL. After 11 weeks of dietary manipulation mice were subjected to an oral glucose tolerance test. (G) chow fed mice and (H) HFD-fed mice and quantified as area under the curve (I). After 11 weeks of dietary manipulation mice were subjected to an insulin tolerance test. (J) chow fed mice and (K) HFD-fed mice and quantified as area under the curve (L). Representative images of ser ^307^ phosphorylation on IRS-1 and normalised to total IRS, ser ^473^ phosphorylation of Akt and normalised to total Akt and ser ^9^ phosphorylation on GSK3β and normalised to total GSK3 β in the liver (M) and skeletal muscle (N) and quantified via densitometry (O and P). Data shown are means ± SEM of n biological replicates. *P< 0.05 **P<0.01 ***P<0.001 ****P<0.0001; a student’s t-test was performed when there were two variables and a one-way ANOVA was performed with Bonferroni post hoc test when there were more than two variables.

We next examined the mice for glycaemic regulation using a fasted oral glucose tolerance test (OGTT) and an insulin tolerance test (ITT). WT mice fed a HFD had a significantly augmented OGTT compared to mice fed a chow diet (Figure 4G/H/I), however, XID mice fed a HFD and subjected to an OGTT display an improved recovery time (Figure 4H/I). WT mice fed a HFD have a significantly augmented ITT compared to mice fed a chow diet (Figure 4J/K/L), no difference was seen in ITT between WT and XID mice fed a HFD (Figure 4K/l).

Having shown that HFD-fed XID mice have improved glycaemic control, we wanted to assess insulin signalling pathways. Serine phosphorylation on IRS-1 in the liver and skeletal muscle is a marker of peripheral insulin resistance ^34^. WT mice fed a HFD had significantly increased Ser^307^ phosphorylation on IRS-1, which leads to a significant decreased phosphorylation of Ser^473^ on Akt and Ser ^9^ on GSK3 β in the liver (Figure 4M/O) and skeletal muscle (Figure 4N/P). When compared to WT mice fed a HFD, XID mice fed a HFD have significantly less Ser^307^ phosphorylation on IRS-1, which attenuated the decrease in phosphorylation of Ser^473^ on Akt and Ser ^9^ on GSK 3 β in the liver (Figure 4M/O) and skeletal muscle (Figure 4N/P) indicating that XID mice are protected from reduced signalling through the IRS-Akt-GSK3 β pathway following HFD feeding resulting in improved glycaemic regulation.

### Adipose associated macrophages from HFD-fed XID mice have an enhanced M2-like phenotype

HFD feeding of mice leads to an expansion of the adipose tissue and an associated increase in immune cell infiltration ^35^. Compared to WT mice fed a HFD, XID-mice fed a HFD displayed a reduction in the density of crown-like structures in epididymal white adipose tissue (eWAT) (Figure 5A/B). The SVF from eWAT was isolated and gene expression analysis revealed a significant decrease in the levels of *CD68* the pan monocyte/macrophage marker (Figure 5C) and reduced *CCL2* a key monocyte chemoattractant/cytokine in XID mice fed a HFD (Figure 5D). Multicolour flow cytometry was then used to assess the polarisation state of adipose tissue macrophages (ATM) from the SVF. ATM from XID mice fed a HFD, display significantly decreased cell surface expression of MHCII (MFI; Figure 5E/F) and decreased mRNA expression of *nos2* (Figure 5G). Additionally, ATM from XID mice also had increased expression of cell surface marker CD206 (MFI; Figure 5H/I), and increased mRNA levels of *Arg1* (Figure 5J) indicating that ATM from XID mice had reduced M1-like phenotype and have enhanced M2-like phenotype. Indeed, *ex vivo* stimulation of ATM with LPS/IFNγ and IL-4 resulted in an exacerbation of the phenotype; ATM from XID mice displayed failed induction of MHCII (Supplementary Figure 4A/B), while IL-4 stimulated ATM from XID mice displayed enhanced cell surface expression of CD206 (MFI) (Supplementary Figure 4A/C).

**Figure 5:**
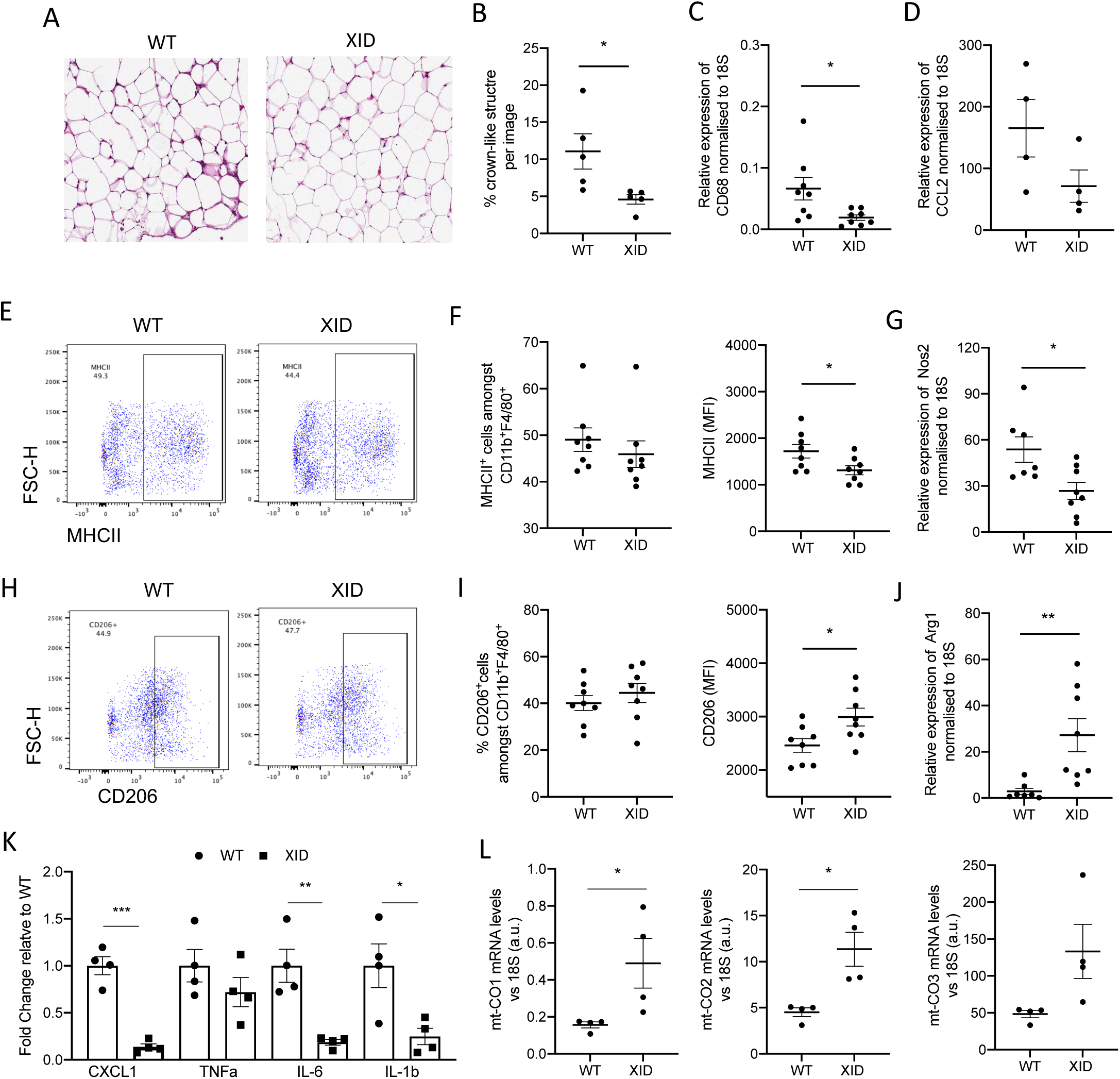
Adipose associated macrophages from HFD-fed XID mice displayed enhanced M2-like phenotype reducing adipose inflammation. WT and XID mice were fed a HFD for 12 weeks and eWAT extract and fixed for staining. (A) Representative images of H&E stained eWAT. (B) quantification of crown-like structures. Relative expression of *Cd68* (C) *and CCL2* (D) in SVF from eWAT isolated from WT and XID mice. (E) Representative flow cytometry plots of adipose associated macrophages (CD11b ^+^ F4/80^+^) from the SVF from eWAT showing expression of MHCII and quantified (F). (G) Relative expression of M1-like marker *nos2* in SVF from eWAT isolated from WT and XID mice. (H) Representative flow cytometry plots of adipose associated macrophages (CD11b ^+^ F4/80^+^) from the SVF from eWAT showing expression of CD206 and quantified (I). (J) Relative expression of M2-like gene *Arg1* in SVF from eWAT isolated from WT and XID mice. (K) Relative expression of pro-inflammatory genes *Cxcl1, tnfa, il-6 and il-1b* in SVF from eWAT isolated from WT and XID mice. (L) Relative expression of mitochondrial complex IV genes *mt-CO1, mt-CO2 and mt-CO3* in SVF from eWAT isolated from WT and XID mice. Data shown are means ± SEM of n biological replicates. *P< 0.05; a student’s t-test was performed when there were two variables, and a one-way ANOVA was performed with Bonferroni post hoc test when there were more than two variables.

Additionally, pro-inflammatory markers (*Cxcl1, Il-6* and *Il-1b*) were reduced in the SVF from XID mice compared to WT mice (Figure 5K). This was also coupled with increased expression of key components of mitochondrially encoded components of electron transport chain Complex IV, *mt-CO1*, *mt-CO2* and *mt-CO3* in the SVF of eWAT from XID mice (Figure 5L). These results show us in the absence BTK signalling macrophages in the eWAT results in reduced M1-like expression profile and enhanced M2-like profile in ATM from HFD fed XID mice, accompanied by increased ability to utilise OxPhos through an upregulation of mitochondrial complex IV genes *in vivo*.

As XID mice are known to have altered B-cell populations we assessed B-cell and myeloid cell populations in the spleen. No difference in the number of CD11b ^+^ myeloid cells was seen in the spleen (Supplementary Figure 5A). As expected XID mice fed HFD had a reduction in the number of CD19 ^+^ B-cells, CD19 ^high^FSC^low^ and CD19 ^+^FSC^high^ B-cells (Supplementary Figure 5B). Suggesting that although systemic B-cells populations are altered there is no effect on myeloid cell number.

### Macrophages from XID mice fed a HFD have enhanced ability to utilise Oxidative Phosphorylation

BMDM from XID mice fed a HFD have enhanced gene expression of classical M2 markers *Arg1* and *Fizz* compared to BMDM from WT mice fed a HFD (Figure 6A). Accordingly, using mRNA expression of electron transport chain Complex IV components, we found that there was significantly increased expression of *mt-CO1*, and a trend towards increased expression in *mt-C02* and *mt-C03* in BMDM from XID mice fed a HFD (Figure 6B). Analysis of protein levels demonstrated there was increased expression of mt-C01 and mt-C02 in macrophages from XID mice fed a HFD compared to WT mice fed a HFD (Figure 6C/D). Furthermore, there was no significant difference in levels of mitochondrial proteins NDUFB8 (from complex I), UQCR2 (from complex III) and ATP5A (from complex V) (Figure 6E/F), consistent with BTK signalling specifically regulating a subset of mitochondrially encoded genes for electron transport chain complex IV.

**Figure 6:**
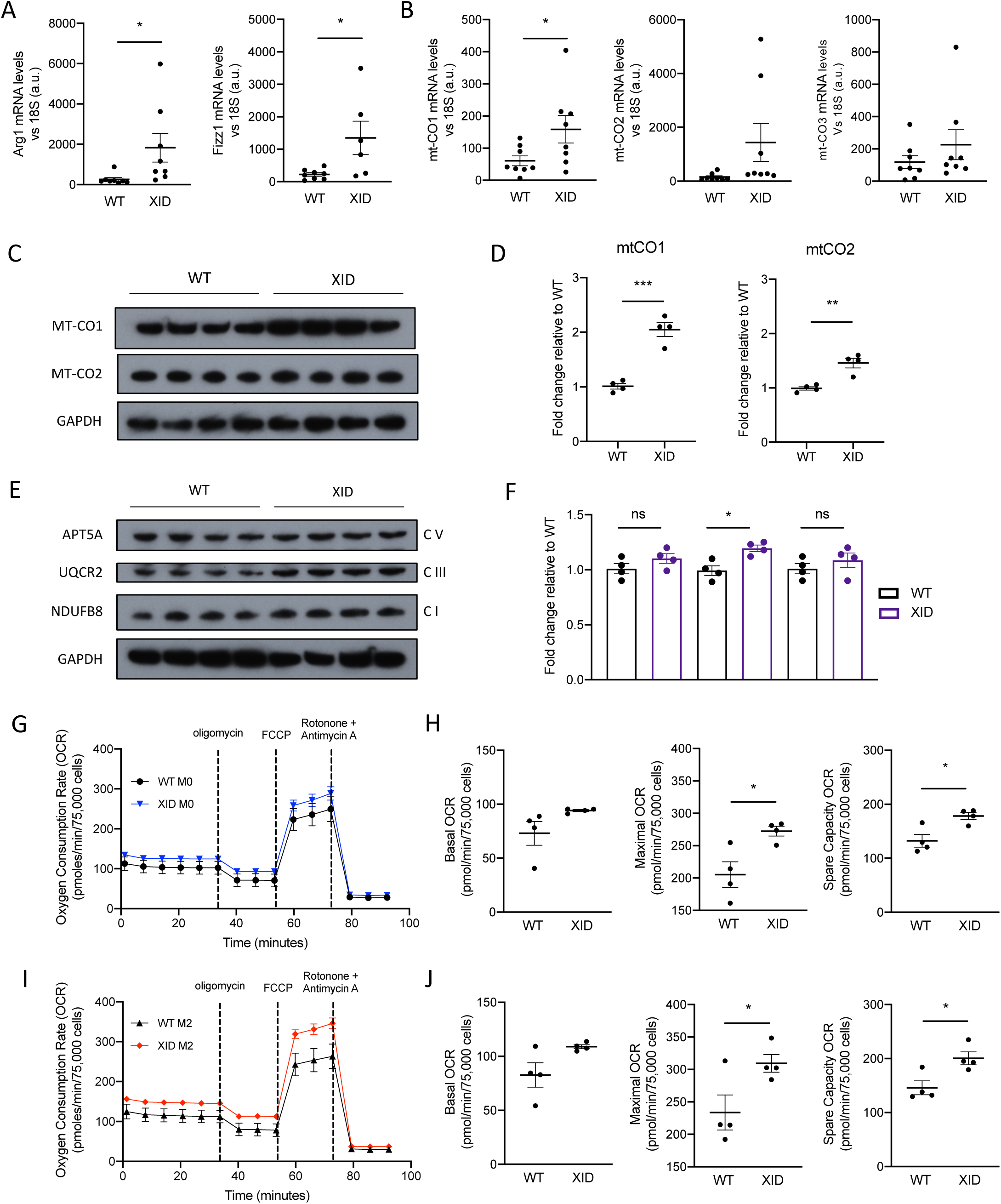
Macrophages from HFD-fed XID mice displayed enhanced ability to utilise oxidative phosphorylation. WT and XID mice were fed a HFD for 12 weeks and bone marrow derived macrophages generated and polarised to M(IL-4). (A) Relative expression of M2-like marker gene *Arg1* and *Fizz1* in M(IL-4) from generated from WT and XID mice. (B) Relative expression of mitochondrial complex IV genes *mt-CO1, mt-CO2* and *mt-C O* i*2*n M(IL-4) from generated from WT and XID mice. (C) Representative western blots of mitochondrial proteins mt-CO1 and mt-CO2 in M0 and M(IL-4) BMDM generated from WT and XID mice. (D) Representative western blots of mitochondrial proteins from Complex I (NDUFB8), Complex II (SDHB), Complex III (UQCR2) and Complex V (APT5A) in M0 and M(IL-4) BMDM generated from WT and XID mice. Oxygen consumption rate (OCR) was measured using XFe96 Seahorse bioanalyzer with compounds used to determine basal, spare capacity and maximum respiration in (G/H) MO BMDM (black solid line -WT; blue solid line - XID) and (I/J) M(IL-4) BMDM (black solid line - WT; red solid line - XID). Data shown are means ± SEM of n biological replicates. *P< 0.05; a student’s t-test was performed when there were two variables, and a one-way ANOVA was performed with Bonferroni post hoc test when there were more than two variables.

Using a Seahorse Analyser to measure OCR we showed that M0 BMDM from XID mice fed a HFD had significantly increased maximal and spare capacity OCR compared to M0 BMDM from WT micef ed HFD (Figure 6G/H), while in IL-4 treated BMDM from XID mice the phenotype was significantly enhanced with a further enhancement in basal OCR, maximal and spare capacity OCR (Figure 6I/J). These data show that inhibition of signalling through BTK allows macrophages to increase expression of key mitochondrial encoded genes which allow for increased ability to utilise OxPhos, which enables enhanced M2-like polarisation.

### Myeloid but not B-cell knock-out of BTK confers a metabolic benefit to mice fed a HFD

To assess which BTK expressing cells were the cellular target in XID mice which protected mice from glycaemic dysfunction we generated both B-cell specific (Figure 7A), and myeloid specific (Figure 7E) BTK knock-out mice, and subjected them to HFD feeding for 12-weeks. When compared wild-type mice, B-cell BTK knock-out (BTK ^fl/y^CD19Cre) mice had no difference in fasted blood glucose level (Figure 7B). When wild-type mice and BTK^fl/y^CD19Cre mice were subjected to an OGTT, there was no difference in response over the 120 min test period (Figure 7C and D). On the other hand, when compared to wild-type mice, myeloid specific BTK knock-out (BTK ^fl/y^LysMCre) mice had significantly lower fasted blood glucose levels (Figure 7F). When wild-type mice and BTK ^fl/y^LysMCre mice were subjected to an OGTT, BTK ^fl/y^LysMCre mice displayed a significantly reduced peak blood glucose spike at 30 mins (Figure 7G), and reduced AUC for the entire 120 min of the test (Figure 7H).

**Figure 7:**
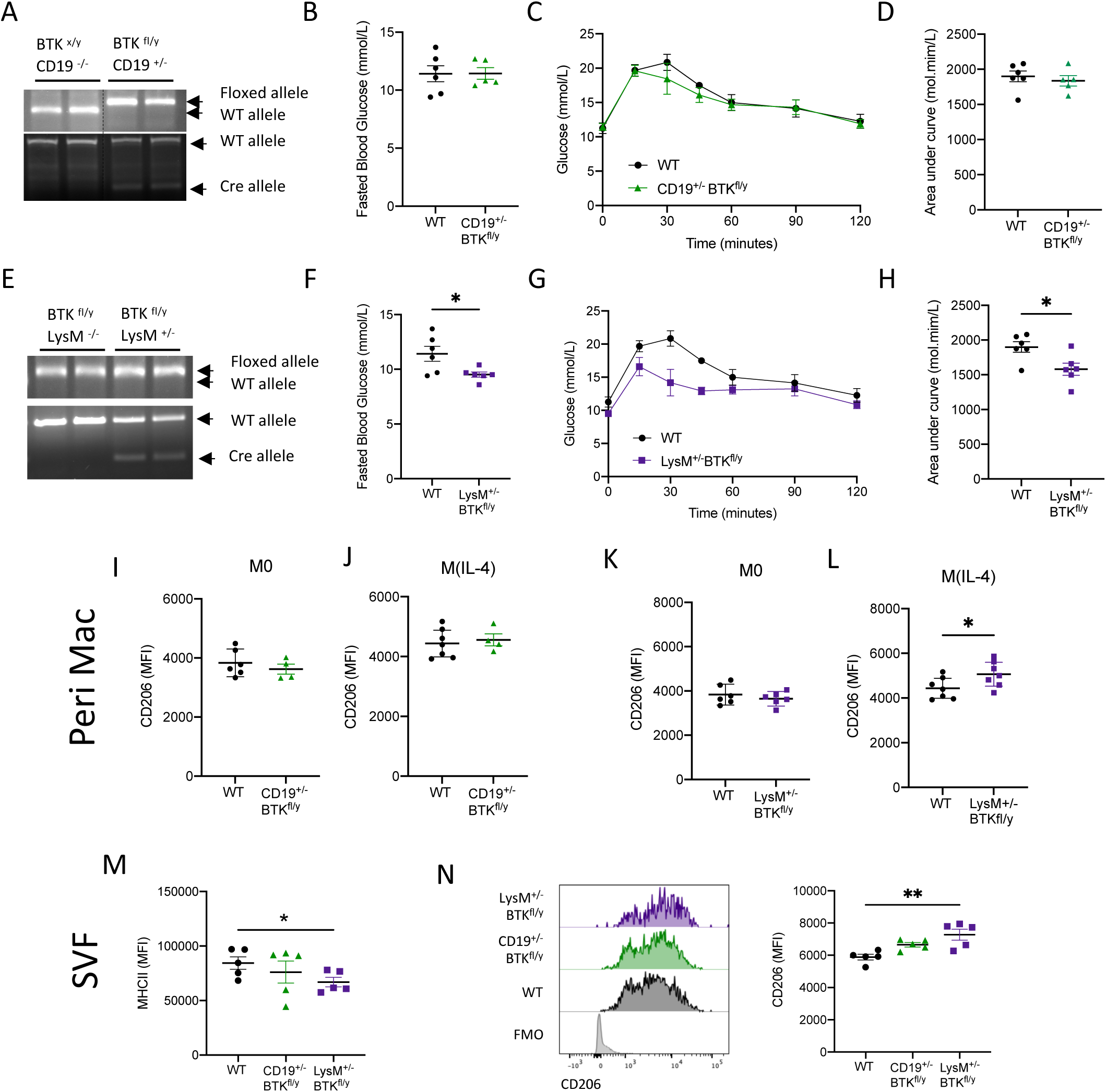
Myeloid but not B-cell specific knock-out of BTK confers metabolic benefit to mice fed a HFD. (A) PCR of genomic DNA in tissue from BTK ^x/y^CD19Cre^-/-^ (WT) mice and BTK ^fl/y^CD19Cre^+/-^ mice. (B) BTK^x/y^CD19Cre^-/-^ mice and BTK ^fl/y^CD19^+/-^ mice were fed a HFD 12 weeks. After 11 weeks of dietary manipulation fast blood glucose was quantified (B). An oral glucose tolerance test performed (C) quantified as area under the curve (D). (E) PCR of genomic DNA in tissue from BTK^fl/y^LysMCre^-/-^ (WT) mice a ^fl^n^/y^LdysMBCrTe^+^K^/-^ mice. (F) ^fl/^B^y^LyTsMK Cre^-/-^ mice and BTK^fl/y^LysMCre^+/-^ mice were fed a HFD 12 weeks. After 11 weeks of dietary manipulation fast blood glucose was quantified (B). An oral glucose tolerance test performed (G) quantified as area under the curve (H). Peritoneal macrophages from BTK ^x/y^CD19Cre^-/-^ mice and BTK^fl/y^CD19^+/-^ were isolated and assessed at baseline (M0) or after polarisation with IL-4. Cell surface expression of M2-like marker CD206 was quantitated in both M0 (I) and M(Il-4) cells (J). Peritoneal macrophages from BTK^fl/y^LysMCre^-/-^ mice and BTK^fl/y^LysM^+/-^ were isolated and assessed at baseline (M0) or after polarisation with IL-4. Cell surface expression of M2-like marker CD206 was quantitated in both M0 and M(Il-4) (I-L). Representative flow cytometry plots of adipose associated macrophages (CD11b^+^ F4/80^+^) from the SVF from eWAT showing expression of MHCII and quantified (M). Representative flow cytometry plots of adipose associated macrophages (CD11b ^+^ F4/80^+^) from the SVF from eWAT showing expression of CD206 and quantified (N). Data shown are means ± SEM of n biological replicates. *P< 0.05 **P< 0.01; a student’s t-test was performed when there were two variables, and a one-way ANOVA was performed with Bonferroni post hoc test when there were more than two variables.

Peritoneal macrophages from BTK ^fl/y^CD19Cre mice display no difference in cell surface expression of M2-like marker CD206 (Figure 7I and J). Peritoneal macrophages from BTK^fl/y^LysMCre display no difference in cell surface expression of CD206 (Figure 7K), however, when polarised to M2 with IL-4, peritoneal macrophages from BTK ^fl/y^LysMCre display increased cells surface expression of CD206 (Figure 7L) compared to wild-type controls. Furthermore, ATM isolated from the SVF of BTK ^fl/y^CD19Cre mice, display no alteration in cells surface of MHCII and CD206 compared to WT mice (Figure 7M). However, ATM from BTK^fl/y^LysMCre mice have significantly increased cell surface expression of CD206 (Figure 7N). Experiments performed using lineage specific BTK knock-out mice clearly demonstrate that BTK signalling is needed to direct macrophage pro-inflammatory signalling leading to glycaemic dysfunction.

### Acalabrutinib treatment of HFD-fed mice improves glycaemic control and enhances ATM M2-like phenotype

We next wanted to test if therapeutic intervention with the BTK inhibitor acalabrutinib could alter ATM polarisation state. C57BL/6J mice were fed a HFD for 6 weeks and then treated with either acalabrutinib (3 mg·kg ^−1^; p.o.; five times per week) or vehicle for a further 6 weeks. Vehicle treated mice on a HFD continue to gain weight while acalabrutinib mice do not gain any further weight (Figure 8A). Mice fed a HFD and treated with acalabrutinib had significantly lower non-fasting blood glucose (Figure 8B), compared with mice fed a HFD and treated with vehicle. When compared to mice fed a HFD diet administered vehicle that underwent an oral glucose tolerance test (OGTT), acalabrutinib treated mice exhibited a shorter elevation in blood glucose levels (Figure 8C), and overall had an improved OGTT assessed by AUC (Figure 8D). These results collectively suggest that mice fed a HFD and treated with acalabrutinib have improved glycaemic regulation.

**Figure 8:**
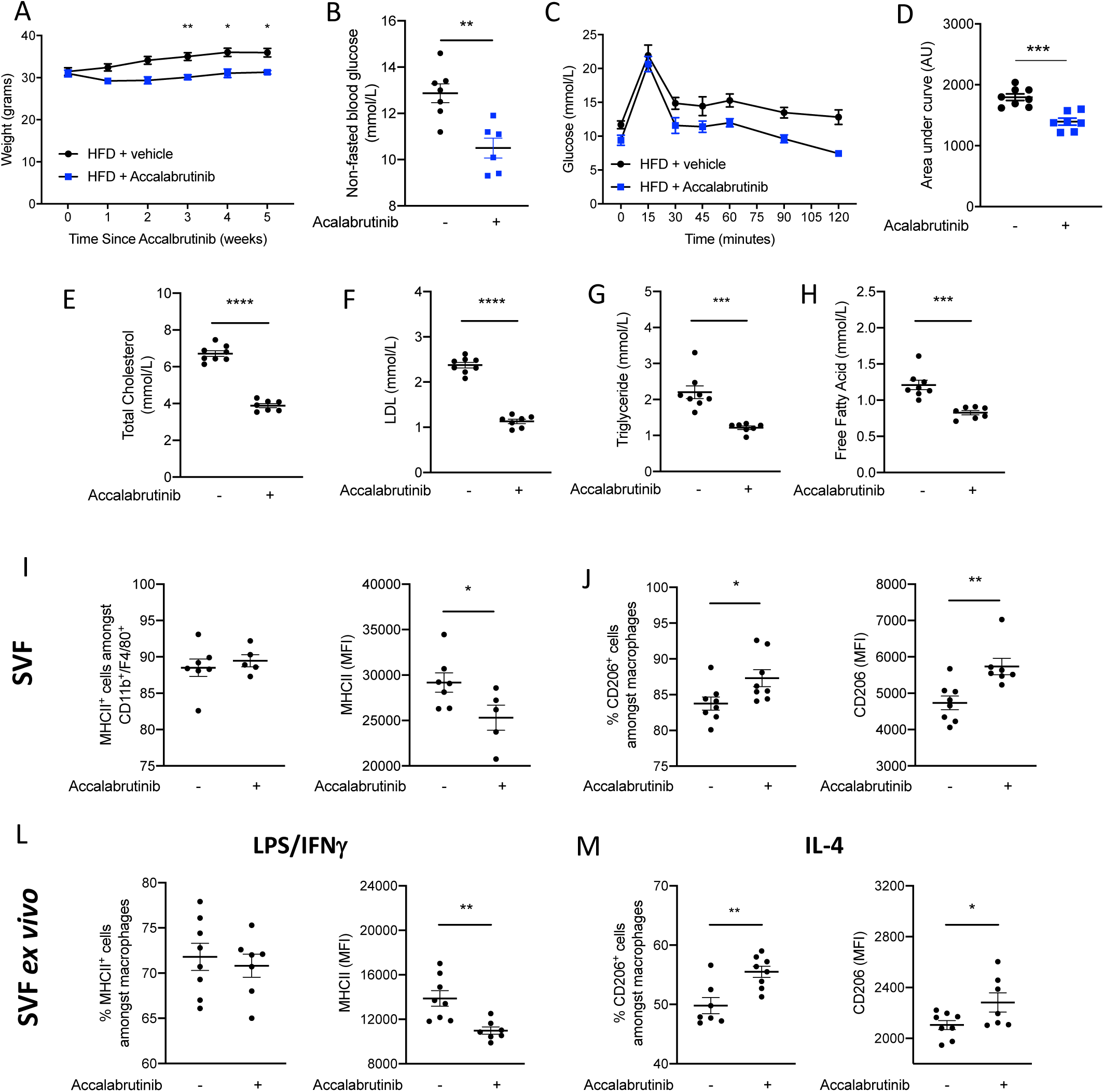
Acalabrutinib treatment to HFD-fed mice improves glycaemic control and enhances adipose associated macrophage M2-like phenotype. WT mice were fed a HFD for 6 weeks then treated with either acalabrutinib or vehicle 5 x per week for a further 6 weeks, body weight was measured each week during dosing (A). Non-fasted blood glucose levels were measured at week 12 (B). After 11 weeks of dietary manipulation mice were subjected to an oral glucose tolerance test (C) and quantified as area under the curve (D). Blood plasma analysis of (E) cholesterol, (F) LDL, (G) triglyceride and (H) free fatty acid. Flow cytometry of adipose associated macrophages (CD11b ^+^ F4/80^+^) from the SVF from eWAT showing expression of MHCII and quantified (I). Flow cytometry of adipose associated macrophages (CD11b ^+^ F4/80^+^) from the SVF from eWAT showing expression of CD206 and quantified at baseline (J). Flow cytometry of adipose associated macrophages (CD11b ^+^ F4/80^+^) from the SVF of eWAT showing expression of MHCII and quantified stimulated with LPS/IFNγ (L). Flow cytometry of adipose associated macrophages (CD11b^+^ F4/80^+^) from the SVF of eWAT showing expression of CD206 and quantified stimulated with IL-4 (M). Data shown are means ± SEM of n biological replicates. *P< 0.05 **P<0.01 ***P<0.001 ****P<0.0001; a student’s t-test was performed when there were two variables, and a one-way ANOVA was performed with Bonferroni post hoc test when there were more than two variables.

Analysis of blood plasma lipid levels revealed that mice fed HFD and treated with acalabrutinib had significantly lower total cholesterol levels (Figure 8E), LDL levels (Figure 8F) decreased TG (Figure 8G) and free fatty acids (Figure 8H) when compared to vehicle treated mice. We next assessed whether pharmacological inhibition of BTK with acalabrutinib could alter ATM polarisation state. No difference in adipose tissue weight was observed in mice treated with acalabrutinib. There was no difference in the number of MHCII+ ATM in mice treated with acalabrutinib, however they had significantly less cell surface expression of MHCII (MFI) (Figure 8I). Importantly, acalabrutinib treatment to HFD-fed mice increased number of CD206+ ATM and significantly increased the cell surface expression of CD206 (MFI) (Figure 8J). Indeed, *ex vivo* stimulation of ATM to M(LPS/IFN γ) and M(IL-4) resulted in an exacerbation of the phenotype. When compared to LPS/IFN γ stimulated ATM from vehicle treated mice, LPS/IFN γ stimulated ATM from acalabrutinib treated mice displayed significantly less cell surface expression of MHCII (MFI) (Figure 8L).

However, when compared to IL-4 stimulated ATM from vehicle treated mice, IL-4 stimulated ATM from acalabrutinib treated mice displayed a further enhancement in cell surface expression of CD206 (MFI) (Figure 8M). Our experiments show that systemic administration of the BTK inhibitor acalabrutinib alters ATM polarisation toward the M2-like pro-resolution phenotype in adipose tissue *in vivo*, which protects mice from glycaemic dysregulation.

### Ablation of macrophages negates the beneficial effects of acalabrutinib treatment on glycaemic regulation in HFD fed mice

To demonstrate that systemic administration of acalabrutinib was inhibiting BTK signalling in macrophages we designed a protocol to deplete macrophages including ATM using clodronate liposomes. After 5 weeks of HFD feeding mice were assigned one of 4 experimental groups; those treated with clodronate liposomes or vehicle (0.5 mg/kg weekly; i.p.), and acalabrutinib (3 mg/kg; 5 x per week; p.o.) or vehicle (Figure 9A). As seen previously, acalabrutinib treatment reduced the number of ATMs compared to vehicle treated mice (Figure 9A/B). Clodronate liposome resulted in a significant decrease in the number of CD11b^+^F4/80^+^ macrophages within SVF isolated from eWAT (Figure 9A/B).

**Figure 9:**
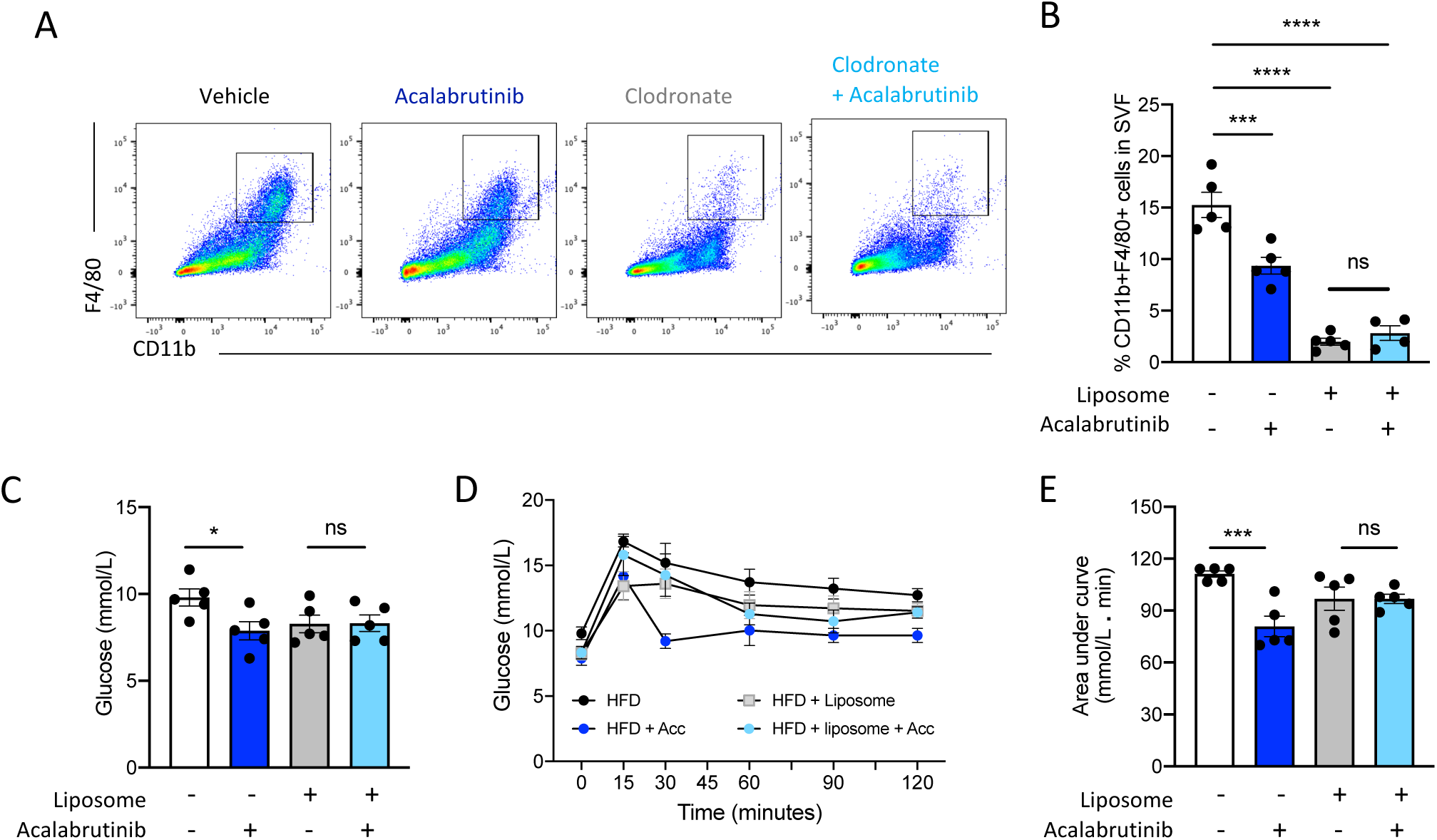
Depletion of ATM negates the beneficial effects of acalabrutinib treatment on glycaemic tolerance in HFD mice. (A) Flow cytometry of adipose associated macrophages (CD11b ^+^ F4/80^+^) from the SVF of eWAT and quantified (B). At week 9 of the protocol fasted blood glucose was measured (C) and mice were subjected to an oral glucose tolerance test (D) and quantified as area under the curve (E). *P< 0.05 ***P<0.001 ****P<0.0001; a one-way ANOVA was performed with Bonferroni post hoc test when there were more than two variables.

At the end of the 10-week protocol mice underwent an OGTT; mice treated with acalabrutinib had a significantly lower fasted blood glucose level (Figure 9C). Compared to mice fed a HFD, administration of clodronate liposomes did not alter fasted blood glucose level (Figure 9C). Importantly, mice co-administered clodronate liposomes to deplete ATM and acalabrutinib displayed no difference in fasted blood glucose level (Figure 9C). Similarly, mice treated with acalabrutinib had a significantly improved OGTT assessed by AUC (Figure 9D/E). Compared to mice fed a HFD, administration of clodronate liposomes did not alter the result of a 120 min OGTT assessed by AUC (Figure 9D/E). Importantly, mice co-administered clodronate liposomes to deplete ATM and acalabrutinib displayed no difference in 120 min OGTT assessed by AUC compared to clodronate liposome treated mice (Figure 9D/E).

## Discussion

In the present study we have shown BTK regulates macrophage polarisation state within murine adipose tissue. Mice with a non-signalling BTK are protected from the immunometabolic consequences of obesity and display improved glycaemic regulation. Using lineage specific knock-out mice we demonstrate unambiguously that myeloid cell, but not B-cell knock out of BTK conferred protection from glycaemic dysfunction in HFD fed mice. Consistent with these findings, systemic therapeutic intervention with the FDA approved BTK inhibitor acalabrutinib protected HFD-fed mice from glycaemic dysregulation. However, the beneficial effects of acalabrutinib treatment were negated in mice deficient in adipose tissue macrophages. With regards to mechanism, inhibition of BTK signalling in macrophages resulted in ATM having a reduced pro-inflammatory and enhanced M2-like macrophage phenotype, reducing both tissue specific and systemic inflammation through upregulation of macrophage OxPhos. This is the first demonstration that using a BTK inhibitor can alter macrophage cellular metabolism and change ATM phenotype *in vivo* . Critically, we have shown that inhibiting BTK signalling in macrophages increased expression of mitochondrial components in Complex I (*mt-CO1*, *mt-CO2* and *mt-CO3*), complex IV (*ND3 and ND4L*) and complex V (*Atp6* and *Atp8*), which allowed macrophages to have enhanced ability to utilise OxPhos.

We have shown that BTK can regulate the expression of mitochondrially encoded genes. Specifically, inhibition of BTK results in increased expression of mtCO1, mtCO2 and mt-CO3, which are integral components of Complex IV which catalyses the transfer of electrons from ferrrocytochrome c to molecular oxygen, allowing ATP to be generated by ATPase synthase^36^. There was also increased expression of two complex I genes, ND3 and ND4L. NF-κB signalling can regulate mitochondrial gene expression ^37^; Cogswell and colleagues found that modulation of NF-κB activation resulted in the loss of expression of both *mt-CO2* and *cytochrome b* ^32^. We and others have previously shown that BTK signalling regulates NF-κB activity in macrophages ^2138^, while in models of obesity NF-κB is activated ^39^ and that inhibition of BTK can reduce NF-κB activity in the same model ^31^. However, we did not detect any p65 protein in the mitochondria of WT or XID BMDM, and inhibiting NF-κB did not result in changes in mtCO1 and mtC02 gene expression levels in our hands, contrary to the findings of Cogswell *et al..* We cannot rule out a non-canonical mechanism of NF-κB which involved gene expression changes independent of p65. We also studied the putative mechanism of BTK:NF-κB regulating mitochondrial gene expression changes in primary BMDM, while previous studies had been conducted in transformed U937 cells.

Macrophages polarisation state is intricately linked with their metabolic status. The switch towards a Warburg-like metabolism in pro-inflammatory macrophages is well established; a shift from oxidative metabolism to glycolytic ATP generation is a result of NO-mediated mitochondrial dysfunction ^9^. Alternatively activated M2/M(IL-4) macrophages, on the other hand, can utilise multiple different sources of energy metabolism, including OxPhos from an intact electron transport chain, glycolysis and ability to use fatty acid oxidation ^10,40,41^. Here we show that macrophages from XID mice have enhanced M2 phenotype, which also have enhanced ability to utilise OxPhos. We propose this is the mechanism by which ATM from XID mice have an enhanced M2 phenotype basally. This phenotype was further enhanced in M(IL-4) macrophage in XID mice. RNA-Seq and protein analysis revealed this was due to an up-regulation of key of mitochondrial complex I and IV components of the electron transport chain. These data demonstrate that BTK signalling can regulate mitochondrial gene expression, but also highlights the dependence of M2-like macrophages on OxPhos for successful M2 polarisation ^42^. To this end, it has been well documented that inhibition of OxPhos using Oligomycin (an F0 subunit ATPase inhibitor) reduces M(IL-4) polarisation ^30,43^. These results demonstrate that cellular metabolism can be altered to enhance macrophage M2-like phenotype by inhibiting BTK signalling *in vivo*.

The development of obesity driven glycaemic dysregulation is associated with the activation of macrophages in the adipose tissue ^44,45,46^; resulting in increased tyrosine phosphorylation of the insulin receptor substrate 1 (IRS-1), reducing PI3K activation which decreases serine phosphorylation of Akt and GSK-3 β and consequently results in reduced insulin signalling ^47,48^. It should be noted that although increased phosphorylation of Ser ^307^ is a marker of insulin resistance in rodents, it is not causative ^34,49^. In our model we see that either genetic or pharmacological inhibition of BTK signalling results in improved glycaemic regulation associated with improved signalling in the IRS-1 complex, therefore allowing for signalling through Akt and GSK-3β ^31^. As insulin resistance and obesity related disorders progress with age; there is a temporal change in the phenotype of macrophages as these metabolic diseases progress. Our analysis of scRNA seq data from the SVF of adipose tissue revealed that homeostatic ‘M0/M2-like’ monocytes/macrophages are present in lean adipose while in obesity there is an increased prevalence of M1-like macrophages. It has been shown that these pro-inflammatory myeloid cells are recruited in CCR2 dependant manner ^5^. We demonstrate that inhibition of BTK signalling not only reduces M1-like ATM, it reduces *CCR2* expression in the SVF but also reduced other pro-inflammatory markers in the SVF from eWAT. We have also demonstrated that inhibition of BTK reduces accumulation of crown-like structures, this could be due to reduced CCR2-dependent recruitment. Inhibition of BTK has been shown to reduced CCR2-dependant monocyte chemotaxis (*in vitro*) and recruitment (*in vivo*) ^50^. BTK signalling in M1-like macrophages in the adipose facilitates the production of a pro-inflammatory microenvironment and secretion of factors that are involved in propagating peripheral insulin resistance for examples IL-6, TNF α and IL-1 β which act in a paracrine manner on insulin sensitive cells resulting in increased inhibitory phosphorylation on IRS-1, reducing Akt signalling which is independent of cell-autonomous BTK signalling.

ATM are believed to contribute to the state of low-grade chronic inflammation that predispose individuals to insulin resistance and the associated metabolic and cardiovascular co-morbidities ^35,51,52^. Phenotypic switching from M1 to M2 in models of diet induced obesity has been demonstrated following dietary intervention with resveratrol ^53^. We have shown that inhibiting BTK with acalabrutinib also induced ATM phenotypic switching towards an M2-like phenotype, suggesting that is it possible to intervene therapeutically and manipulate macrophage phenotype *in situ* and that this can protect against glycaemic dysregulation and improve signalling through the IRS1/Akt/GSK3 β pathways. By depleting ATM and treating HFD mice with acalabrutinib we were able to demonstrate unambiguously that that the beneficial effects of acalabrutinib are mediated through inhibiting macrophage BTK. Macrophage dependant BTK, but not B-cells BTK signalling resulted in macrophage phenotype switching, and resulted in improved glycaemic regulation, highlighting the importance of pro-inflammatory signalling from macrophages in the development of obesity induced metabolic syndrome.

Acalabrutinib is a second-generation EMA/FDA approved BTK inhibitor that is highly selective but there are currently there are numerous other BTK inhibitors in clinical trials which are also highly selective. Having these potential therapeutics available makes BTK a tractable target for drug repurposing to treat the low-grade chronic inflammation associated with diabetes. It should be noted that acalabrutinib treatment to HFD-fed mice resulted in a small but significant reduction in body weight which could influence systemic metabolism. However, when mice were treated with clodronate liposomes to deplete macrophages the beneficial effect of acalabrutinib treatment was abrogated. Therefore, we propose that the beneficial effects in mice treated with acalabrutinib are drug specific and not due to the reduced weight of drug treated animals. Additionally, the benefit in glycaemic regulation in acalabrutinib mice is consistent with the phenotype seen in Xid mice and LysMCre BTK^fl/y^ mice fed a HFD.

Currently, all licensed BTK inhibitors have been used to target B-cell activation in chronic disease. B-cells have been shown to produce diabetes associated antibodies in both type-1 and type-2 diabetes, however the incidence of diabetes associated antibodies is much lower in type-2 diabetes. Kendell *et al* . have shown that Btk-deficient (BTK ^-/-^/NOD mice) have reduced incidence of diabetes while administration of anti-insulin IgH could restore diabetic incidence ^54^. However, how relevant anti-insulin antibodies are in our model of diet induced obesity is not known, as mice remain partially insulin sensitive. Indeed, we did not see a benefit in glycaemic regulation in B-cell specific knock out of BTK. Nishimura *et al.* have shown that regulatory B-cells in the subcutaneous fat and eWAT produce high levels of IL-10, which is needed to exert homeostatic functions and supress adipose inflammation. XID mice have been shown to have increased CD19 ^low^ B-cell numbers which produce IL-10, which could act in a paracrine manner to increase macrophage M2-like polarisation ^55^.

Further studies are needed to fully elucidate if there is cell to cell cross talk between B-cell and macrophages in eWAT, and if BTK signalling is involved. Our work points to an important role for BTK signalling in macrophages in the development of glycaemic dysregulation in diabetes, with less contribution of B-cell expressed BTK, as the beneficial effects of BTK inhibition are lost when macrophages are ablated in eWAT. Although we have shown that macrophages from XID have a cell autonomous mechanism to increase OxPhos and enhance their capacity to polarise towards an M2-like state, this does not rule out extrinsic signalling from other cell types that regulate macrophage cellular functions.

In conclusion, we have demonstrated that BTK signalling regulates macrophage M2-like polarisation state by altering mitochondrial encoded gene expression allowing for an enhanced rate of OxPhos. Critically, BTK expression is elevated in adipose tissue macrophages from obese patients, while key mitochondrial gene (mtCO1, mtCO2 and mtCO3) are decreased in inflammatory myeloid cells in obese patients. Inhibition of BTK signalling, genetically, or therapeutically protects HFD-fed mice from developing glycaemic dysregulation by improving signalling through the IRS3/AKT/GSK3B pathway. Importantly it also induces a phenotypic switch in adipose tissue macrophages from a pro-inflammatory M1-state to pro-resolution M2-like phenotype, by rewiring macrophage metabolism towards OxPhos. This reduces both adipose tissue specific and systemic inflammation and protects mice from the immunometabolic consequences of obesity. Therefore, in BTK we have identified a macrophage specific, druggable target that can regulate ATM polarisation by altering cellular metabolism.

## Author contribution and acknowledgements

GSDP and DRG conceptualised the study. GSDP, MC, ADvD, MC and GE did the experimental work and analysed the data. GSDP drafted the manuscript. MS and YN recruited healthy subjects and facilitated blood sampling. MC and GE conducted western blotting of murine tissue samples. SP recruited XLA patients and facilitated blood sampling. GSDP, MC, ADvD, GE, YN, MS, SYP, CT KMC and DRG reviewed and edited the manuscript. This work was funded by the British Heart Foundation Grant Number: RG/15/10/23915 to DRG and RG/15/10/31485, RG/17/10/32859 and CH/16/1/32013 to KMC. National Institute for Health Research (NIHR) Oxford Biomedical Research Centre (RG/13/1/301810) to OXAMI Study. MS was supported by the Alison Brading Memorial Scholarship in Medical Sciences, Lady Margret Hall, University of Oxford. Pump Prime award from the Oxford British Heart Foundation Research Excellence: Grant Number: RE/13/1/30181 to GSDP and DRG. Novo Nordisk-Oxford Postdoctoral Fellowship to ADvD.

The transcriptomic analysis was supported by the Wellcome Trust Core Award Grant Number 203141/Z/16/Z with additional support from the National Institute for Health Research (NIHR) Oxford Biomedical Research Centre. The views expressed are those of the author(s) and not necessarily those of the NHS, the NIHR or the Department of Health. We thank the Oxford Genomics Centre at the Wellcome Centre for Human Genetics (funded by Wellcome Trust grant reference (203141/Z/16/Z) for the generation and initial processing of the sequencing data.

We thank Dr Wasif Khan for the kind donation of CD19-Cre BTK fl/fl mice. No conflict of interest needs to be declared.

Prof David R Greaves is the guarantor for this work, as such, had full access to data in the study and takes responsibility for the integrity of the data and the accuracy of the data analysis.

**Supplementary Figure 1: Phosphorylation of BTK in polarised human macrophages.**

(A) Representative immunoblots for phosphorylation of Tyr ^223^ on BTK and total BTK of protein lysates from THP-1 cells stimulated with LPS/IFNγ, IL-4 or vehicle.

**Supplementary Figure 2: Phosphorylation of BTK in polarised human macrophages.**

(A) Top 10 pathways computed by Pathway enrichment analysis of using the top 76 differentially expressed genes between M(IL-4) BMDM generated from WT and XID mice.

**Supplementary Figure 3: NF-kB does not regulate mitochondrial gene expression.**

(A) WT or XID BMDM were stimulated with IL-4 or vehicle then stained with MitoTracker DeepRed (red), p65 (green) followed by counter stained with Hoechst stain (blue). Images were captured under oil immersion at X63 zoom. (B) Relative expression of mitochondrially encoded genes *mtCO1* and *mtCO2*, and nuclear encoded *Cybb* in THP-cells stimulated with LPS/IFNγ, IL - 4 or vehicle with or without IKK - 16 (1 µ M). (C) WT or XI DBMDM we restimulated with IL-4 or vehicle then stained with MitoTracker DeepRed (red) and quantified. (D) Relative expression of mitochondrially encoded genes *mtCO1, mtCO2* and *mtCO3 in* BMDM stimulated with LPS/IFNγ, IL-4 or vehicle. *P< 0.05; a student’s t-test was performed when there were two variables, and a one-way ANOVA was performed with Bonferroni post hoc test when there were more than two variables.

**Supplementary Figure 4: Adipose associated macrophages polarised ex vivo from HFD-fed XID mice displayed enhanced M2-like phenotype.**

(A) Representative flow cytometry plots from of adipose associated macrophages (CD11b ^+^ F4/80^+^) from the SVF from eWAT stimulated ex vivo with LPγ,SI/LI-F4Nor vehicle. (B) Flow cytometry of adipose associated macrophages (CD11b^+^ F4/80^+^) showing expression of MHCII and quantified following LPS/IFN γ stimulation (B). Flow cytometry of adipose associated macrophages (CD11b^+^ F4/80^+^) from the SVF from eWAT showing expression of CD206 and quantified following IL-4 stimulation (C). Data shown are means ± SEM of n biological replicates. *P< 0.05; a one-way ANOVA was performed with Bonferroni post hoc test when there were more than two variables.

**Supplementary Figure 5: XID mice have reduced B-cells and normal myeloid cell counts.**

(A) Representative flow cytometry plots of myeloid cell (CD11b ^+^) f r o m t h e s p l e e n a n d quantified. (B) Representative flow cytometry plots of B-cell population (CD19 ^+^; CD19 ^low^ CD19^high^FSC^low^ and CD 1^+^F9SC^high^) from the spleen and quantified. *P< 0.05 **P< 0.01; a student’s t-test was performed when there were two variables, and a one-way ANOVA was performed with Bonferroni post hoc test when there were more than two variables.

