## Supplementary Material for "OxPhos in adipose tissue macrophages regulated by BTK enhances their M2-like phenotype and confers a systemic immunometabolic benefit in obesity"

**Supplementary Figure 1:** Phosphorylation of BTK in polarised human macrophages.

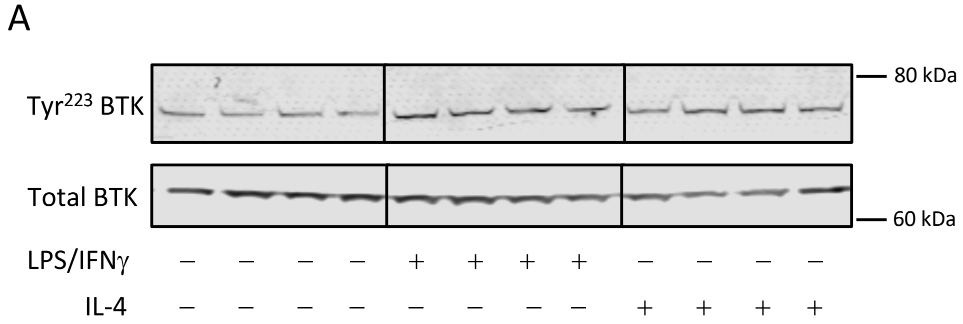


**Supplementary Figure 2:** Ingenuity Pathway Analysis (IPA) of the DE genes between WT and XID M(IL-4) macrophages.

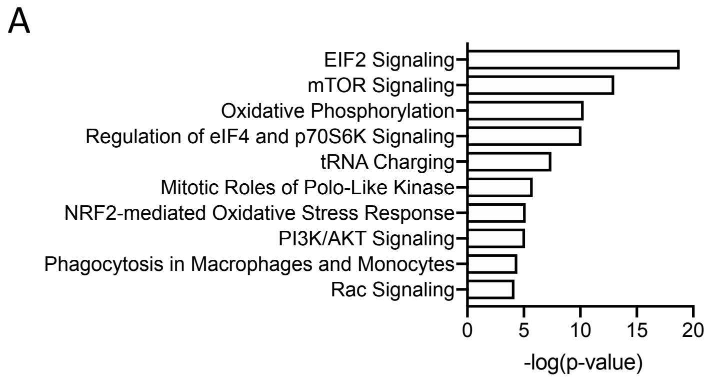

**Supplementary Figure 3:** NF-kB does not regulate mitochondrial gene expression.

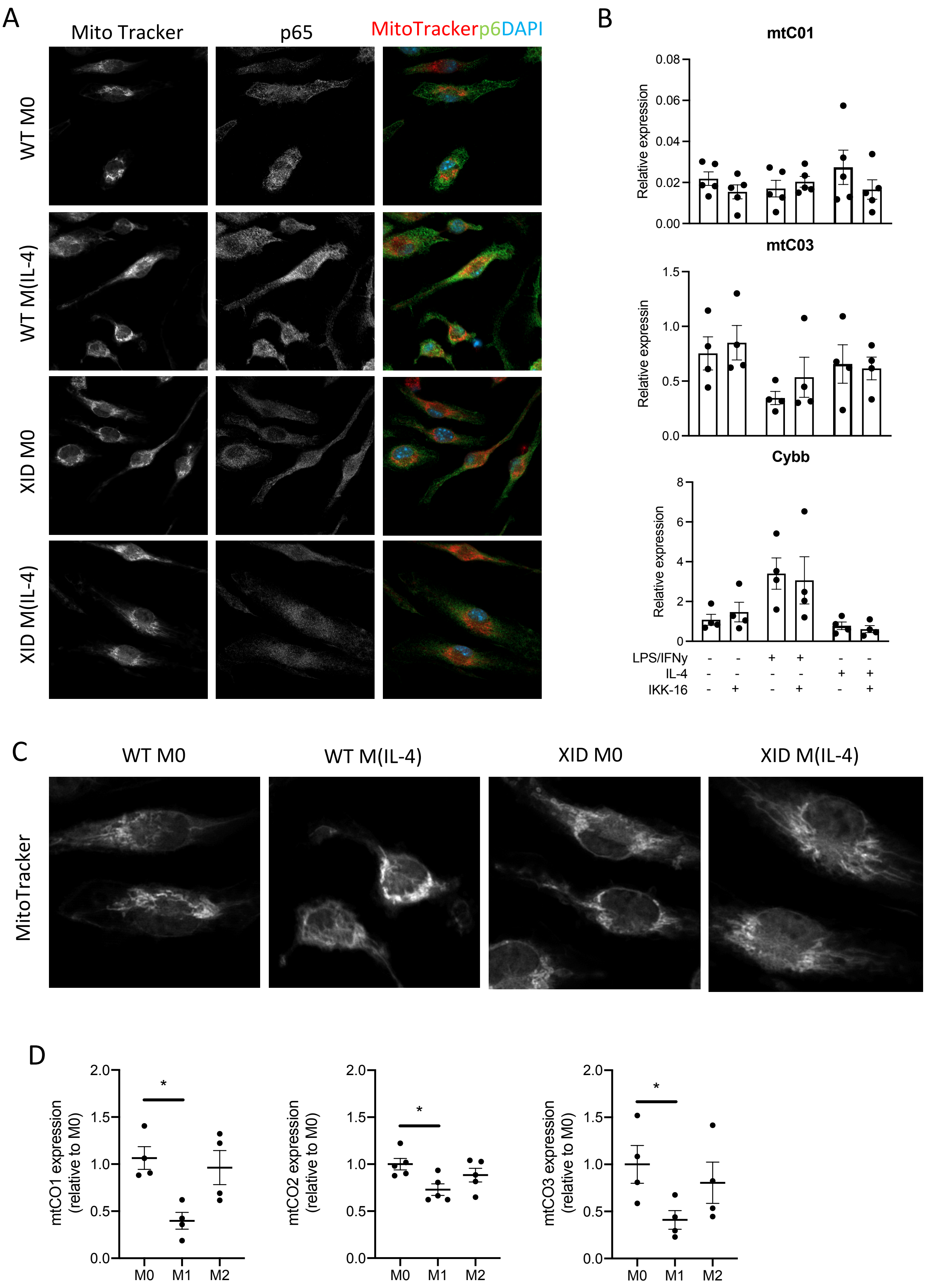

**Supplementary Figure 4:** Adipose associated macrophages polarized *ex vivo* from HFD-fed XID mice displayed enhanced M2-like phenotype

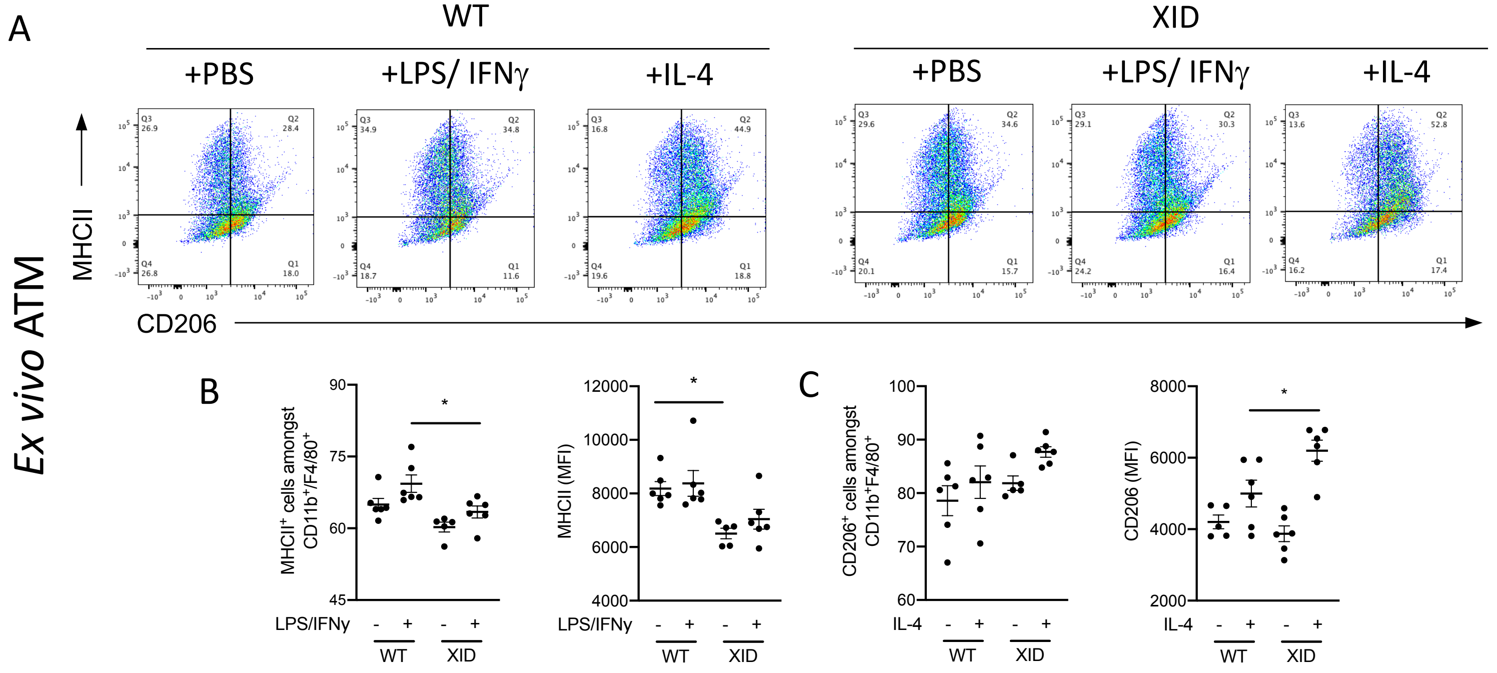

**Supplementary Figure 5:** XID mice have reduced mature B-cells and normal myeloid cell counts.

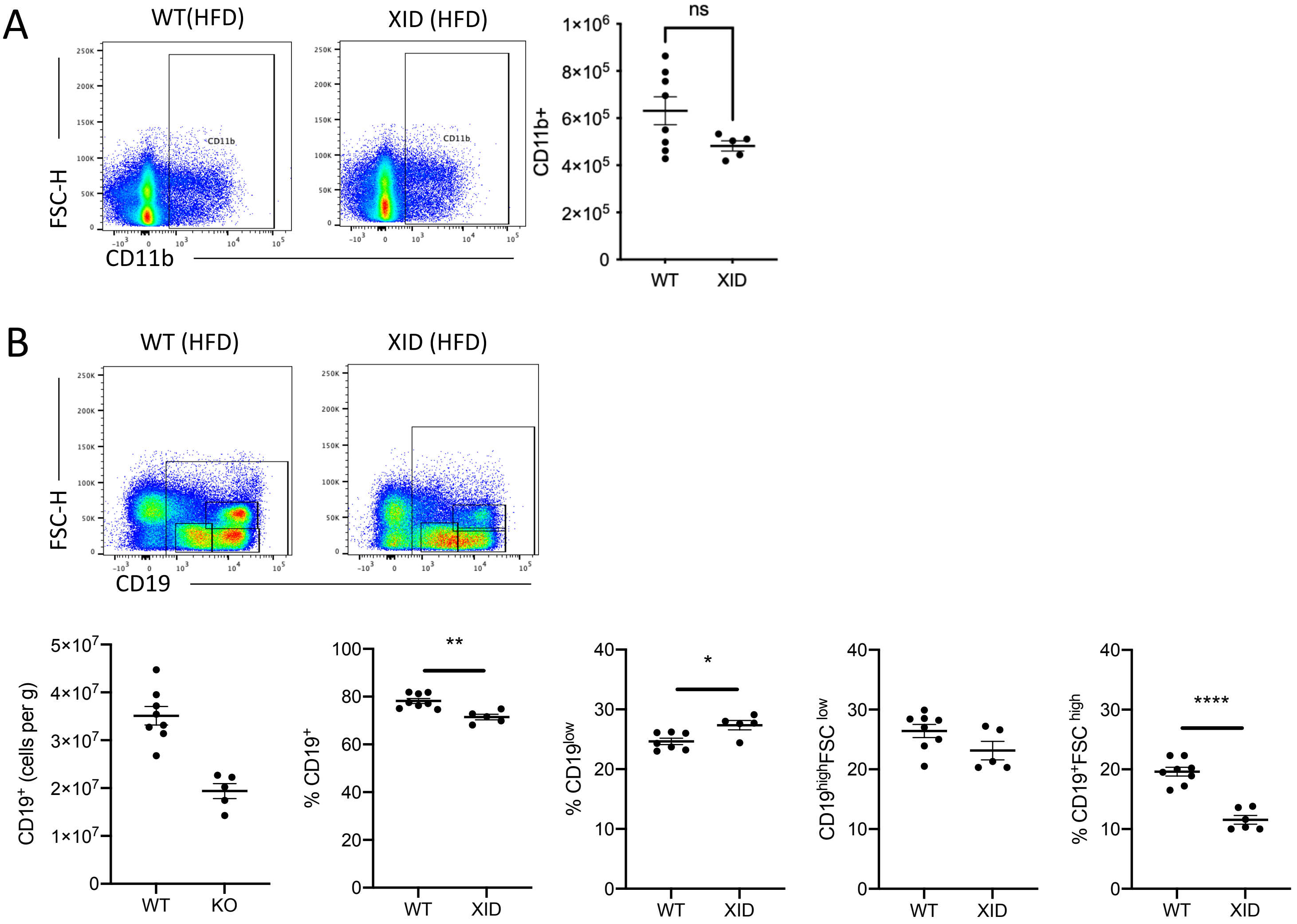
